# Dystrophin is a mechanical tension modulator

**DOI:** 10.1101/2022.12.23.521750

**Authors:** Arne D. Hofemeier, Mariam Ristau, Mattias Luber, Till M. Muenker, Fabian Herkenrath, Bruno Schmelz, Lisa-Marie Scharfenstein, Matthias Brandt, Polina Malova, Mina Shahriyari, Malte Rinn, Daniel Haertter, Malte Tiburcy, Wolfram-H. Zimmermann, Christof Lenz, Kamel Mamchaoui, Anne Bigot, Jo Nguyen, Penney M. Gilbert, Clémence Lièvre, Jean-Baptiste Dupont, Philip Barrett, David Mack, Timo Betz

**Author notes:** Contributing authors.

## Abstract

Duchenne muscular dystrophy (DMD) represents the most common inherited muscular disease, where progressing muscle weakness leads to loss of ambulation and premature death. DMD is caused by mutations in the dystrophin gene, and is known to reduce the contractile capacity of muscle tissue both *in vivo*, and in reconstituted systems *in vitro*. However, these observations are based on mechanical studies that focused on stimulated contractions of skeletal muscle tissue. Seemingly paradoxical, upon evaluating bioengineered skeletal muscles produced from DMD patient derived myoblasts we observe an increase in myosin motor mediated homeostatic tissue tension that strongly correlates with decreased stimulated tissue strength, suggesting the involvement of dystrophin in regulating the baseline homeostatic tension level of tissue. This was confirmed by three independent pairs of isogenic cell lines, one of each lacking dystrophin compared to the isogenic dystrophin-expressing control. From this we speculate that the protective function of dystrophin also supports cellular fitness via active participation in the mechanosensation to achieve and sustain an ideal level of tissue tension. Hence, this study reveals fundamental novel insights into skeletal muscle biomechanics and into a new key mechanical aspect of DMD pathogenesis, provided by increased homeostatic tissue tension.

## 1 Introduction

Skeletal muscle is one of the most abundant tissues in the human body and its proper function is crucial for existential functions such as voluntary movement, thermogenesis, breathing and maintaining posture [1–3]. Dysregulation contributes to life-limiting genetic muscle disorders such as Duchenne muscular dystrophy (DMD), affecting 1 in 3500 - 5000 male live births [4], being the most common inherited muscular disease. Even though the causative gene *DMD* and its associated protein, dystrophin, is known and studied since the 1980s [5, 6], DMD still is an incurable disease. Eventually this devastating disorder becomes lethal within the third decade of a patients’ life due to cardio-respiratory failure [7].

Although affected boys appear clinically normal at birth, muscle biopsies reveal necrotic muscle fibers, tissue inflammation and fibrosis on a microscopic level even before the onset of typical symptoms [7, 8]. Despite the commonality of a genetic dystrophin loss, an intriguing finding is that muscles decline in form and function at different paces when comparing different muscle groups [9]. Characteristically, patients with DMD develop difficulties walking during childhood and later become wheelchair dependent in their teens [10]. As the disease progresses, DMD patients further lose upper body function leading to respiratory distress and cardiomyopathy [10, 11]. Taken together, the pathological severity of DMD not only varies widely depending on the age of the patient but also on the muscle being evaluated [9, 12].

In recent years, DMD research has led to the implementation of a variety of therapeutic approaches, e.g. corticosteroid anti-inflammatory drugs and exon skipping strategies. Unfortunately, these therapies only delay disease progression to a certain extent [13]. Gene therapy clinical trials are underway and offer a promising therapeutic approach [14]. However, the efficacy and long-term impact of such therapies in humans remains unknown. As such, more knowledge is urgently needed to fully understand the development and progression of DMD from the molecular to the cellular and the system levels.

The protein impacted in DMD, dystrophin, is one of the largest intracellular proteins that links the actin cytoskeleton to the transmembrane dystroglycan protein complex. This triggered the hypothesis that it stabilizes the muscle cell membrane against forces by stretching like a spring during contraction [6]. The large number of spectrin repeats in the dystophin protein supports this idea as they can act as shock absorbers. Under high tension, folded spectrin repeats are known to unfold [15], thereby essentially relaxing the transmitted tension. Interestingly, the opening of spectrin repeats can also function as a mechanosensing mechanism by exposing cryptic binding sites. This was demonstrated in the case of *α*-actinin, another actin binding protein [16]. Besides these tension buffering, or potentially tension sensing capacities, dystrophin is proposed to be essential for force transmission between the acto-myosin contractile machinery and the sarcolemma, including its surrounding extracellular matrix (ECM) [17, 18]. Loss of dystrophin leads to destabilization of the muscle fiber membranes which results in elevated intracellular calcium concentrations as well as reactive oxygen species (ROS) levels. Additionally, it is accompanied by the absence of neuronal nitric oxide synthase (nNOS) at the sarcolemma and reduced overall nitric oxide (NO) levels [19–21]. Although state of the art research can draw from multifactorial and complex experimental results, a coherent picture merging the many roles of dystrophin is still to be defined. Partially, this is due to missing genetic models that actualize the human DMD phenotype sufficiently. For instance, the sheer size of the dystrophin gene with 2.2 Mbp does not allow for simple replacement by mutants. Additionally, the most commonly used *mdx* mouse strain fails to recapitulate important phenotypic hallmarks observed in humans, such as the loss of ambulatory ability [22].

For this reason, numerous *in vitro* strategies have emerged in recent years for cultivating self-organizing, human multinucleated myotubes and also other cell types within 3D extracellular matrix (ECM) scaffolds [23–27]. Based on this, the first 3D bioengineered human skeletal muscle disease models were reported only recently, and offer great opportunities to further fundamental research and drug development [28–33]. By contrast to 2D myotube culture, these approaches support studies of the force generating capacity of normal and DMD muscle tissue upon stimulating a contraction. However, in contrast to recent results showing that the constant tissue forces and tension are highly relevant in tissue development [25, 34, 35], studies on muscle functionality mainly focused on stimulated force generation, including DMD [31]. In doing so, a key parameter is less well studied; the homeostatic (e.g. baseline) tension of the tissue that is present even in the non-stimulated situation at rest and is actively maintained. Tensional homeostasis is a well studied mechanism in 2D cell cultures [36, 37], and tensile forces are known to be crucial for nuclei migration during fiber maturation and symmetry breaking [26, 38]. Additionally, a recent study on zebrafish and mice embryonic development showed that this homeostatic tension is directing polarity of the forming muscle, making it a key parameter for correct muscle development [39]. Therefore, the homeostatic tension is distinct from the commonly reported passive resting tension, which is affiliated to elastic forces in the connective tissue fraction of muscle and also to viscoelastic myofibrillar structures (e.g. titin)[40–42]. As the resting tension is experimentally identified with the external load that allows the largest contractile force generation, the homeostatic tissue tension is furthermore a distinct quantity from a measurement point of view and can be accessed *in vivo* by analyzing the retraction velocity of an muscle after cutting.

Even though tissues are known to actively regulate their tensional homeostasis [43, 44] and the dystrophin binding partner nNOS was reported to play a crucial role in regulating mechanical loading in skeletal muscle [18, 45, 46] and relaxation of smooth muscles [47], the homeostatic tissue tension within skeletal muscle has not been in focus in the context of DMD, quite surprisingly. In closing this knowledge gap, our study reveals a surprising, and initially counter-intuitive relationship, whereby strongly contracting healthy muscle tissue are antagonized by low homeostatic tension, while DMD muscle tissue show a higher homeostatic tension but offer limited contractile function in response to electrical stimulation. This was confirmed by three pairs of isogenic cell lines. First, healthy iPSC as well as healthy immortalized myoblasts were genetically engineered to DMD knockout lines leading to elevated homeostatic tissue tension and decreased stimulated contractility independent of different 3D matrices (fibrin vs. collagen) used. Additionally, we found in another iPSC-derived cell system [48], that the presence of a malfunctioning dystrophin protein in tissues showed increased homeostatic tension and decreased stimulated contraction when compared to tissues derived from the same cell line that had a recovered functional dystrophin protein by CRISPR/Cas9 mediated exon skipping. Furthermore, *in vivo* measurements of homeostatic tension in healthy and mdx mice by muscle cutting experiments further demonstrated an increased homeostatic tension if dystrophin is absent. These findings support the central role of dystrophin for the tensional homeostasis of muscles.

## 2 Results and Discussion

### 2.1 Myotube and muscle tissue morphology is dependent on dystrophin

Skeletal muscle precursor cells, also called ‘myoblasts’, can be isolated from patients’ biopsies and immortalized for further long-term applications [49]. Here, we used cell lines derived from 8 different donors and from biopsies originating from three different muscle groups (Table 1). As recently reported, these cell lines are capable of forming functional biomimetic skeletal muscle tissue within 3D protein scaffolds by self-organizing around two vertical posts in custom engineered devices [30, 31]. With these human immortalized myoblast cell lines from healthy and DMD backgrounds, we cultured bioengineered skeletal muscles within a 3D fibrin-Geltrex™ scaffold over two weeks (Figure 1 a,c). For that purpose, we utilized our previously described platform for high resolution imaging [26], that enables the cultivation of bioengineered tissue between two hanging flexible posts in closest proximity to the thin glass bottom (Figure 1 b,d). The platform is milled from polymethyl methacrylate (PMMA) as a modular two part system. The myoblasts within the protein scaffold are seeded into the ellipsoid wells of the bottom part. The lid contains a pair of posts for each well. All of the human immortalized cell lines self-organized to form tissues around the two hanging posts (Figure 1 e). Precise measurement of the tissue width revealed that dystrophic tissues are on average thinner when compared to healthy tissues (Figure 1 g). The resulting averages are dominated by drastically reduced dystrophic tissue widths of bioengineered skeletal muscles derived from cells that originated from leg muscle biopsies (Figure 1 h). By contrast, bioengineered dystrophic skeletal muscle tissues produced using cells originating from paravertebral biopsies are slightly thicker. Additionally, when investigating the myotubes formed within the bioengineered skeletal muscles at two weeks of differentiation, additional morphological differences become apparent between the healthy and dystrophic tissues (Figure 1 f). Although immunohistological staining revealed multinucleated myotube formation in all models, in general the dystrophic tissues using cells derived from leg muscle biopsies exhibit immature and thinner myotubes with less sarcomere structures aligning in parallel.

**Fig. 1.**
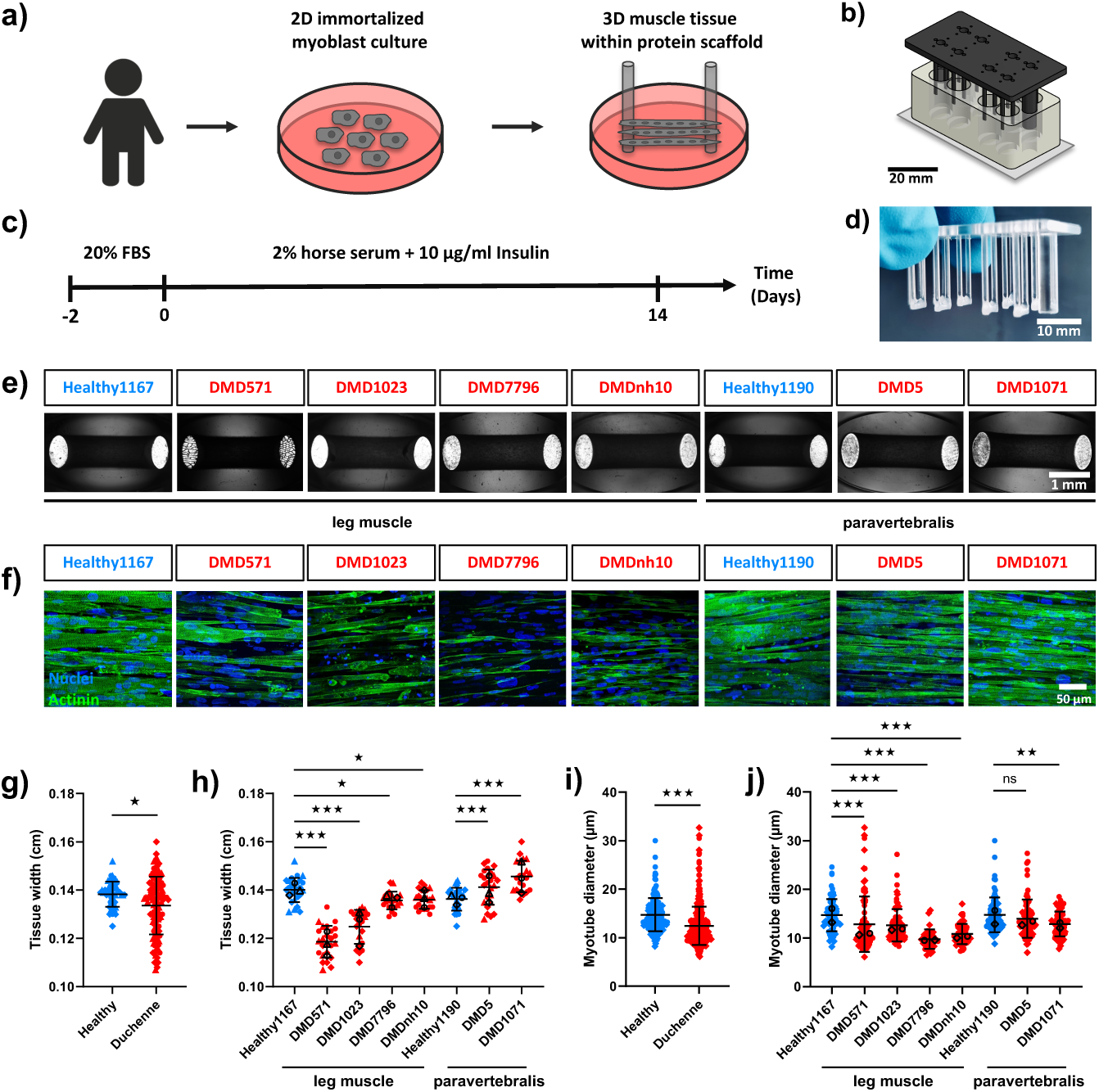
Tissue appearance and myotube diameter size evaluation of bioengineered dystrophic human skeletal muscles. a) Schematic overview of 3D bioengineered skeletal muscle tissue generation within a custom PMMA culture mold (b). c) Timeline of 3D human skeletal muscle tissue cultivation *in vitro*. d) Representative capture of two weeks old human bioengineered skeletal muscle tissue anchored to the tips of the PMMA posts. e) Representative brightfield images of two weeks old healthy and DMD human skeletal muscle tissue *in vitro*. f) Representative flattened confocal stacks of multinucleated myotubes within healthy and DMD human skeletal muscle tissue immunostained for sarcomeric-alpha-actinin (green) and nuclei counterstained with DRAQ7 (blue). g-h) Tissue width of healthy and DMD human skeletal muscle tissue, pooled (g, n=51/135 tissues per group from N=3 independent experiments each) and separated between the different patients (h, n=27/25/22/20/22/24/25/21 tissues per group from N=3 independent experiments each). i-j) Myotube diameter of healthy and DMD human skeletal muscle tissue, pooled (i, n=158/388 myotubes per group from N=2 tissue from two independent experiments each) and separated between the different patients (j, n=77/82/80/35/56/81/76/59 myotubes per group from N=2 tissue from two independent experiments each). All results are presented as mean *±* SD. Statistical differences in g) and i) were analysed by unpaired t-test, h) and j) by one way ANOVA multiple comparison, with p *<* 0.05 considered as significant. Independent experiments performed on different days are represented by different symbols, whereof the median of each independent experiment is presented in black.

**Table 1.** Human immortalized patient derived myoblasts used in this study.

| cell line name | phenotype | mutation in DMD | origin | age |
| --- | --- | --- | --- | --- |
| Healthy1167 | healthy | none | fascia-lata | 20 |
| Healthy1190 | healthy | none | paravertebral | 16 |
| DMD571 | Duchenne | Deletion exon 52 | fascia-lata | 10 |
| DMD1023 | Duchenne | Stopp codon exon 59 | quadriceps | 11 |
| DMD7796 | Duchenne | Deletion exon 42-43 | quadriceps | 4 |
| DMDnh10 | Duchenne | Deletion exon 44-50 | quadriceps | 2 |
| DMD5 | Duchenne | Duplication exon 2 | paravertebral | 14 |
| DMD1071 | Duchenne | Deletion exon 45-52 | paravertebral | 13 |

Overall, the dystrophic myotubes were smaller in width compared to healthy myotubes (Figure 1 i). One exceptional case was myotubes within tissues produced using the paravertebral derived DMD5 cell line, where only insignificant differences were observed (Figure 1 j). DMD5 has an exon 2 duplication which is susceptible to revertant fiber formation due to an alternative translation initiation site in exon 6 [50], a phenotype we observe in tissues produced using this line (Supplement 1a). These data are in exact compliance with previous reports of Ebrahimi et al. [31], who very recently found revertant fiber formation in DMD5 bioengineered muscles for the first time. In sum, these results suggest that the cell line biopsy tissue origin can be reflected by phenotypes observed *in vitro* in bioengineered tissues, and are consistent with clinical results concluding that different muscle groups are affected by DMD with different severity [9]. Despite these findings, at the current stage the clinical relevance of the observed differences requires more detailed experiments in close interaction with patients. Therefore, we focus in the following on the cell-biological and biophysical aspects that can be studied using these immortalized cell lines.

### 2.2 Stimulated contractility versus homeostatic tissue tension

Since the human bioengineered dystrophic skeletal muscles showed gross morphological differences when compared against healthy tissues, we next investigated their functional contractility. Here, we sought to distinguish between the two types of contractile forces that the bioengineered muscles exert on the posts. First, the stimulated contractility that is widely used to examine muscle strength upon a stimulus, e.g. electrical pulses. Second, the unstimulated contractile force of the bioengineered tissue that to this point has not been in the focus of the field, and which we term ‘homeostatic tissue tension’. Homeostatic tissue tension can be determined by analysis of the post deflection that naturally arises during tissue development without any external stimulus. It is important to note that the homeostatic tension is mediated by active contractile forces cells exert on their environment, and not an outcome of ECM compaction. We therefore distinguish the homeostatic tension from the passive resting tension that is related to connective tissue stiffness and elastic myofibrillar proteins [40–42].

To study the stimulated tissue contractility, we positioned electrodes exactly behind each vertical post and electrically stimulated the bioengineered skeletal muscles with a pulsed electrical field of 5 V at 20 Hz to investigate full tetanus strength (Figure 2 a, Supplement 2). Since these electrodes tightly fit into the holes present on the lid of the chambers, we can assure a very similar position in every experiment. During stimulated contraction, healthy as well as dystrophic skeletal muscles exerted sufficient forces to deflect the posts (Figure 2 b, Supplement 3). Using the spring constant of the posts (39 *µN/µm*) reported in our previous studies [26], we calculated the contractile forces from post deflection videos (example given in Figure 2 c) of healthy and dystrophic skeletal muscles upon electrical stimulation. As expected, healthy bioengineered skeletal muscles typically exerted more force on the posts (∼0.2 ± 0.1 mN) during a 20 Hz stimulated tetanus contraction when compared to tissues from a dystrophic background, which exerted around 0.1 ± 0.1 mN in average (Figure 2 e). When comparing the different donor-derived skeletal muscle tissues separately, we found that this phenotype was more pronounced in the tissues produced using myoblasts from a leg muscle group background (Figure 2 f). The Healthy1167 cell line elicited almost 0.2 ± 0.1 mN stimulated contractile force. However, DMD571, DMD1023, DMD7796 and DMDnh10 did not even reach 0.05 ± 0.02 mN on average. Again, the paravertebral derived DMD5 cell line breaks ranks with a stimulated contractile force of more than 0.2 ± 0.09 mN on average that is similar to the paired healthy sample. Paravertebral derived DMD1071 was significantly weaker than the paravertebral derived Healthy1190. Taken together, these data are consistent with previous reports [31], which stresses the reliability of our system. Interestingly, the stimulated contractile forces of the human immortalized myoblast derived tissues strongly correlated with the myotube diameter (Supplement 4). This observation may support functional estimations of the biomimetic skeletal muscle strength from morphological data in the future.

**Fig. 2.**
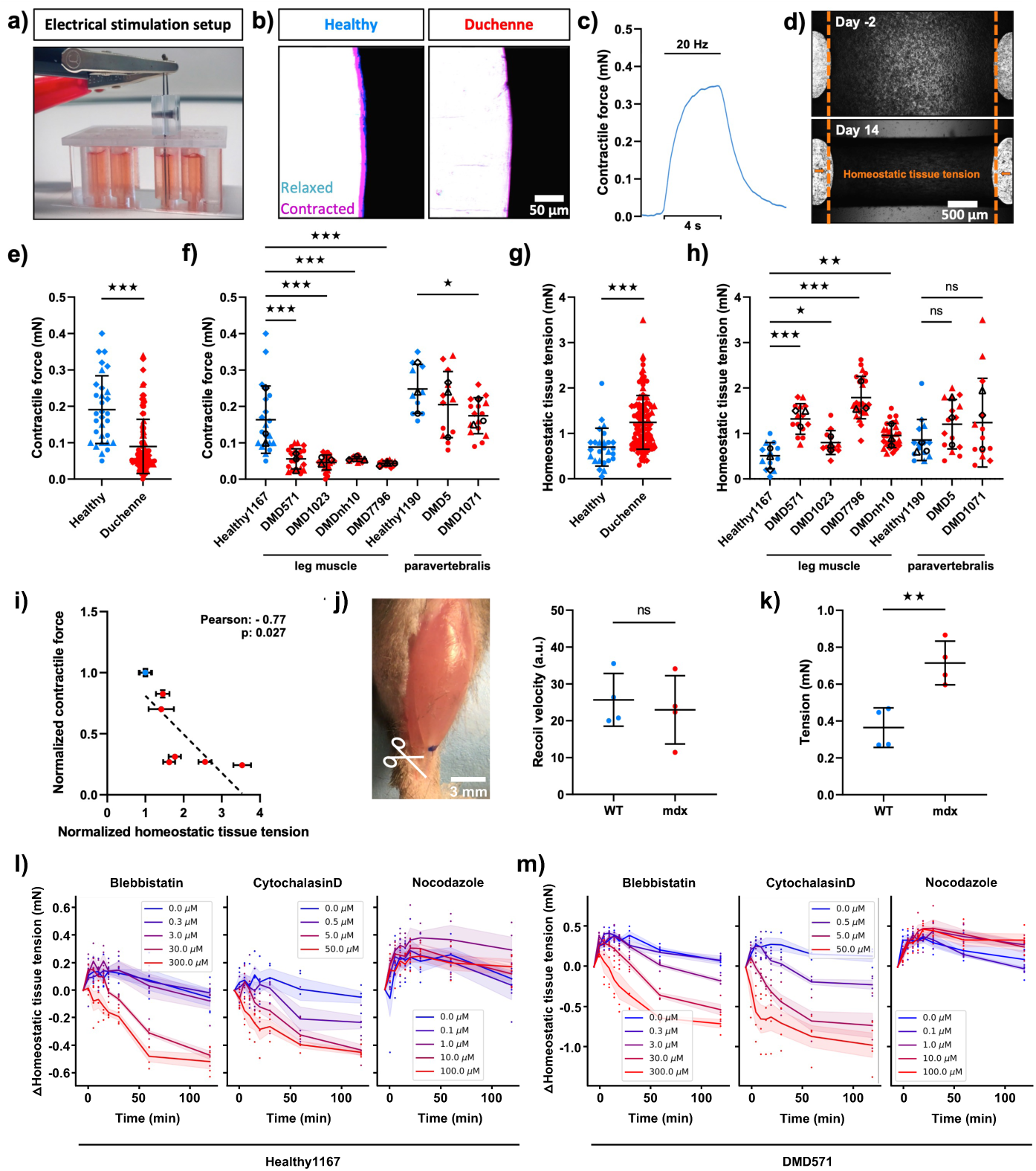
Quantification of human bioengineered dystrophic skeletal muscles’ contractile forces and homeostatic tissue tension. a) Setup for electrical stimulation of human skeletal muscles *in vitro*. b) Post deflection by healthy and DMD human bioengineered skeletal muscles within the PMMA culture mold upon 20 Hz electrical stimulation. c) Representative contraction curve recorded by post deflection analysis upon a 20 Hz electrical stimulation. d) Representative brigthfield images acquired after tissue seeding and after two weeks of differentiation used for quantification of the homeostatic tissue tension via post deflection analysis. e-f) Contractile force of healthy and DMD human skeletal muscle tissues upon 20 Hz electrical stimulation, pooled (e, n=31/98 tissues per group from N=3 independent experiments each) and separated between the different patients (f, n=21/21/21/12/13/10/12/16 tissues per group from N=3 independent experiments each). g-h) Homeostatic tissue tension of healthy and DMD human bioengineered skeletal muscle tissues, pooled (g, n=26/107 tissues per group from N=3 independent experiments each) and separated between the different patients (h, n=12/14/12/22/24/14/15/13 tissues per group from N=3 independent experiments each). i) Anti-correlation between the normalized contractile force and homeostatic tissue tension of healthy (blue) and DMD (red) bioengineered muscles. As we normalize the leg and paravertebralis muscles by the respective healthy tissue, the two healthy values collapse on a single value (blue) at 1. j-k) Recoil velocity (j) and resulting tension (k) analysis of tibialis anterior (TA) muscle from WT and mdx mice (n=4/4 TA muscle per group from N=2/2 mice). l-m) Dose dependent (purple to red color scheme) homeostatic tissue tension decrease upon Blebbistatin, CytochalasinD and Nocodazole treatment on Healthy1167 (l, n=4 tissues per condition from N=2 independent experiments) and DMD571 (m, n=4 tissues per condition from N=2 independent experiments) compared to DMSO-treated control tissues (blue). Results in i), l) and m) are presented as mean *±* SEM, all other results as mean *±* SD. Statistical differences in f) and h) were analysed by one way ANOVA multiple comparison, all others by unpaired t-test, with p *<* 0.05 considered as significant. Independent experiments performed on different days are represented by different symbols, whereof the median of each independent experiment is presented in black.

Besides the stimulated force, we wondered if the homeostatic tension, which represents a continuous, unstimulated force generated by the muscle cells is also affected in DMD derived tissue. The importance of this tension that is actively generated by the cells has been demonstrated during development, where the homeostatic tension controls muscle orientation and thereby proper muscle development [39]. To obtain the homeostatic tension, we used 2 complementary approaches. First, we measured post relaxation during tissue disassembly after addition of SDS to 14d old tissues. The post stiffness was then used to obtain the homeostatic tension. Furthermore, we recorded the post distance shortly after seeding the cells (day −2) and compared this distance with the post distance after differentiation and muscle tissue formation (day 14) as shown in Figure 2 d. The homeostatic tissue tension was obtained by multiplying the shortening of the post distance between these two timepoints with the spring constant of the posts (39 *µN/µm*, [26]).

Against intuition, we observed an increased homeostatic tension in muscle tissue originating from DMD patients in comparison with tissue from healthy donors. This results in a surprising inverse relationship between the level of stimulated contractility and the measured homeostatic tissue tension (Figure 2 i). Specifically, dystrophic bioengineered muscles showed an overall higher homeostatic tissue tension (1.2 ± 0.6 mN) compared to the healthy controls (0.7 ± 0.4 mN) (Figure 2 g). Again, this phenotype is dominated by the significant differences observed from cell lines derived from leg muscle biopsies (Figure 2 h) wherein the homeostatic tissue tension was increased two to threefold in DMD compared with the healthy bioengineered muscles. Indeed, the homeostatic tissue tension is the dominant force exerted on the vertical post as it is more than three times higher on average than the stimulated tetanus contraction of healthy bioengineered muscles. For dystrophic bioengineered muscles, the homeostatic tissue tension was more than ten times higher than the stimulated contractile force emphasizing the severe imbalance of these two mechanical properties. In addition, higher homeostatic tissue tension was in most cases accompanied by thinner tissue width (Figure 1 h). Given the increased homeostatic tension, the decreased width can be easily explained by an overall higher cellular contractility during tissue formation, where also a higher contraction in the direction perpendicular to the post-post direction leads to thinner tissues. Similar to a thinning of a stretched elastic, that becomes thinner if placed under higher tension. Although the tissue widths differed between the groups, the specific homeostatic tissue tension and the specific contractile force (Supplement 5 a,b), normalized to the cross sectional areas, yielded in similar observations of decreased specific force and increased specific homeostatic tension of DMD tissue. Here, also the paravertebralis derived DMD tissues showed decreased specific contractile force. To test if the ECM composition was relevant for the found increase in homeostatic tension in DMD cells, we confirmed the overall result in tissues generated in a collagen I ECM matrix (Supplement 5 c,d).

Next we wondered if there was a correlation between the stimulated and homeostatic force generation. To test this, we normalized the stimulated and homeostatic forces to the healthy tissue controls of leg and paravertebralis muscle and analyzed them in a scatter plot. This analysis revealed a strong anticorrelation between homeostatic tissue tension and the contractile force generation (Figure 2 i). In simple words, the reduced stimulated contractility of diseased muscles is accompanied by a relative increase in homeostatic tension. This suggests an interesting new hypothesis, namely that the poor stimulated contractility of DMD derived bioengineered muscles is due to the elevated homeostatic tissue tension that leads to an incapability of the tensed tissues to contract further. To extend this point, it is as if the diseased muscle is wasting a large fraction of finite force generation capacity in the unstimulated situation. Interestingly, the increased homeostatic tension may also contribute to the common clinical reports of stiffened Duchenne skeletal muscles [51], which was concluded to be the effect of progressing fibrosis. As it is known *in vitro* that increased cellular contractility leads to higher cell stiffness [52], the increased homeostatic tension might also increase tissue stiffness without the requirement of increased fibrosis. Since the myoblast cell lines are clonally derived [49], without any fibroblasts or other cell types, we hence speculate that increased homeostatic tissue tension in dystrophic skeletal muscles may contribute to higher muscle stiffness at rest (also see Supplement 6 a,b) additionally to fibrosis as reported before [53]. To determine the stiffness of the reconstituted muscle tissue, we directly tested stiffness by applying a known force to a reconstituted tissue and recorded the resulting strain. Supplement 6b shows that indeed the DMD derived tissues are not only more tensed, but with a Young’s modulus of 3.8 ± 2 kPa and 7 ± 4 kPa, also stiffer than the tissue derived from the healthy patient (1.4 ± 0.6 kPa). This suggests that the increased homeostatic tension in reconstituted skeletal muscle tissue contributes to stiffening similar to other non-muscle tissues [54]. Aiming at accessing the homeostatic tissue tension also *in vivo*, we compared the recoil velocities of the tibialis anterior (TA) muscle from WT and mdx mice upon cutting the TA tendon (Figure 2 j), which did not reveal differences. To obtain the tension from the recoil velocity, the viscosity of the tissue needs to be considered. The higher the viscosity, the slower the recoil velocity in case of similar homeostatic tension. Since the mdx TA muscle was very recently shown *in vivo* to have indeed a higher viscosity [55], the same recoil velocity in mdx TA muscle compared to healthy TA corresponds to an increased tension of mdx TA muscle. Using reported average TA cross sectional area [56] and length [57], we indeed obtain a significantly higher tension in mdx TA muscle of 0.71 ± 0.12 mN (Figure 2 k) in comparison to WT TA (0.36 ± 0.11 mN) which is in excellent agreement with our observations in bioengineered microtissues (Figure 2 g-h). To further rule out that the homeostatic tissue tension is a matrix-driven passive phenomenon, rather then actively driven and regulated by the muscle cells, we inhibited in a dose dependent manner myosin2 motors pharmacologically in Healthy1167 bioengineered microtissue using Blebbistatin (Figure 2 l). Indeed, we find that the homeostatic tissue tension was decreased by around 0.5 ± 0.1 mN within 120 min for the highest concentration (300 *µ*M). Since inhibition of uncapped actin polymerization using CytochalasinD led to a decrease of 0.45 ± 0.04 mN in the highest concentration of 50 *µ*M (Figure 2 l) we can speculate that most of the tensional homeostasis is driven by cortical acto-myosin contraction, interestingly. Contrary, Nocodazole treatment did not lead to a decrease in homeostatic tissue tension, strengthening that hypothesis. This is again in great compliance with a recent study suggesting that the static tension in bioengineered skeletal muscle is mediated by the undifferentiated myoblast portion within a C2C12 generated tissue [58]. To ensure that the effect of the drugs is a systematic effect across cell lines and cell origin, we repeated these experiments first in leg derived DMD571 bioengineered microtissue (Figure 2 m) and in 3 different paravertebralis derived myoblast cell lines using the highest dose (Supplement 6 c), all confirming that the homeostatic tension is driven by active cell contraction and not corresponding to passive tension. Intrigued by the overall variability of our results across the eight different, patient derived cell lines, we wondered if the elevated homeostatic tissue tension in dystrophic bioengineered skeletal muscles is a tissue phenomenon or the outcome of already mechanically ill-equipped single cells.

### 2.3 TFM reveals increased average traction forces of DMD myotubes and myoblasts

To compare the homeostatic force generation properties at the tissue level to those at the myotube and even single myoblast level, we exploit the fact that both mechanically pull on their environment, which can be quantified by Traction Force Microscopy (TFM) [59–62]. Here we use state of the art Bayesian estimation of regularization parameters [63] to ensure proper regularization. TFM uses elastic polyacrylamide (PAA) gels that include fluorescent nanopar-ticles to track potential deformations caused by the adherent cells that are pulling on the gels (Figure 3 a).

**Fig. 3.**
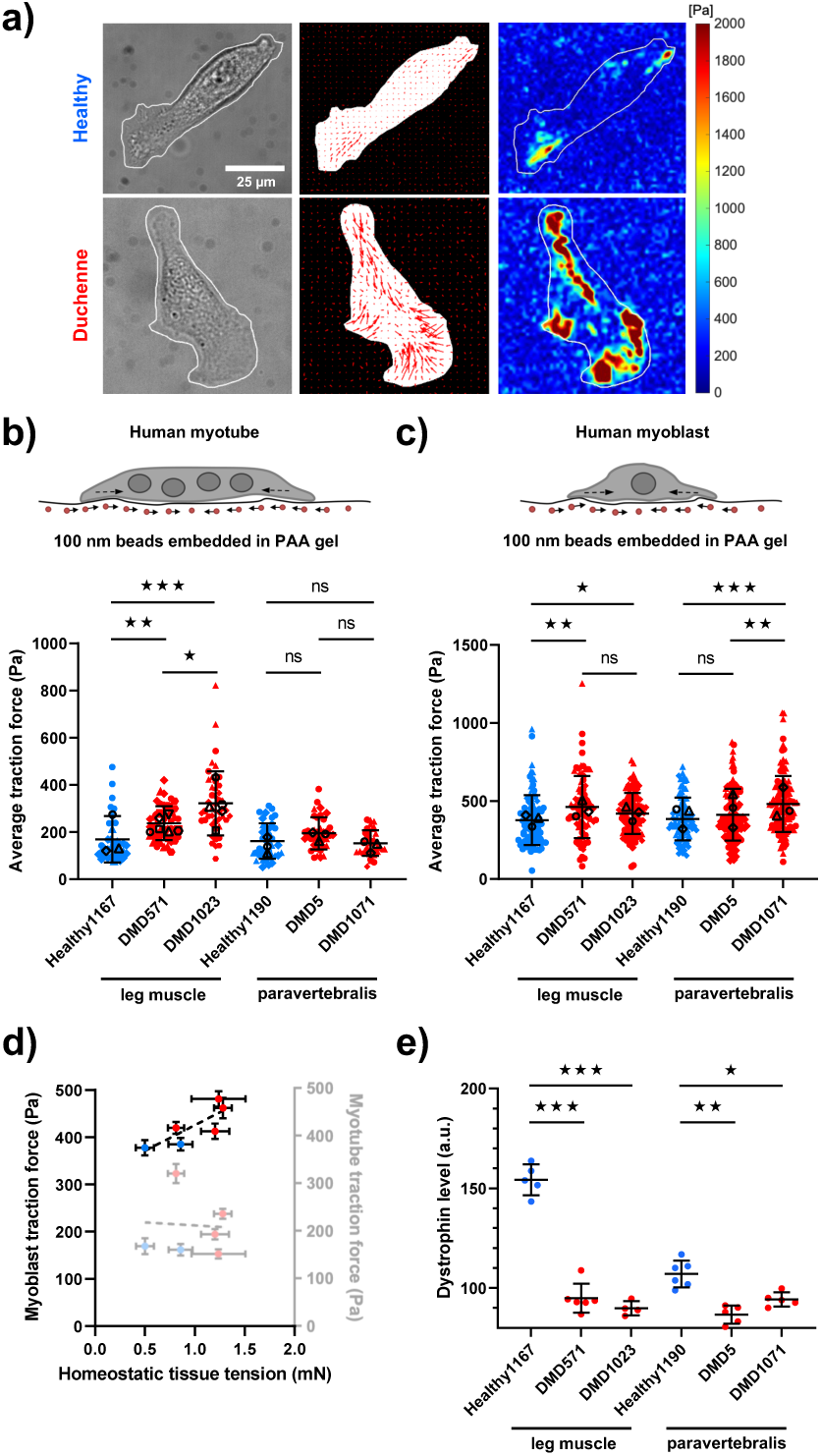
Homeostatic contractility of DMD myotubes single myoblasts. a) Representative cell mask, vector map and force map of a healthy and DMD myoblast obtained during traction force microscopy (TFM) analysis. b-c) Schematic TFM setup sketch and collected average traction forces of healthy and DMD myotubes (b, n=34/46/46/38/38/33 myotubes per group from N=3/6/5/3/3/3 independent experiments) and single myoblasts (c, n=94/81/108/89/103/119 myoblasts per group from N=3 independent experiments each). d) Correlation between the average traction forces of myoblasts/myotubes and homeostatic tissue tension of healthy (blue) and DMD (red) bioengineered muscles. e) Changes in dystrophin levels of healthy and DMD myoblasts determined by data independent acquisition mass spectrometry (DIA-MS) (n=5/6/4/6/5/5 samples per group collected from N=1 independent experiment). Results in d) are presented as mean *±* SEM, all other results as mean *±* SD. Statistical differences were analysed by one way ANOVA multiple comparison with p *<* 0.05 considered as significant. Independent experiments performed on different days are represented by different symbols, whereof the median of each independent experiment is presented in black.

We first performed TFM of myotubes differentiated for one week on 36 kPa (Young’s modulus) PAA gels. By analysing the bead displacements within the PAA substrate, we obtained average traction forces of leg-derived DMD myotubes that were also significantly higher than the healthy control (Figure 3 b). For paravertebralis derived myotubes we did not find significant differences in average traction forces. This confirms in a 2D *in vitro* assay the increased homeostatic tissue tension of leg-derived dystrophic bioengineered muscles.

Next, we wondered if the undifferentiated mononucleated Duchenne single myoblast exhibit this altered contractile behaviour. Therefore, we performed TFM of undifferentiated single myoblasts on physiological 15 kPa (Young’s modulus) PAA gels that closer resemble the tissue properties in the 3D chamber system. Comparing the human immortalized myoblasts from healthy and DMD patients, we found that, on average, single DMD myoblasts pulled more strongly on their culture substrates than did the healthy cells (Figure 3 c), which was reflected by an increase in traction forces, and also by elevated strain energy and contractile moments (Supplement 8). Hence, already at the single DMD myoblast level, the increased contractility reflects a higher homeostatic tension, which is also found in dystrophic myotubes and the dystrophic bioengineered muscles.

To further characterize the extent to which the 3D homeostatic tension of bioengineered muscle tissue can be explained by the single cell contractility, we looked at the overall correlation between these two quantities. As expected from the former results, the traction forces of the myoblasts strongly correlate with the homeostatic tension on a tissue level (Figure 3 d). This suggests that the elevated dystrophic tissue tension is not a collective tissue effect but can be explained, at least in part, by an increased contractility on the single myoblast level. This increased contractility, however, cannot be attributed to elevated Myosin2 levels (Supplement 7a), which suggests a signaling reason. This interpretation reinforces, that the homeostatic tension of a tissue is, in contrast to the passive resting tension, actively regulated by the cell via signaling cascades similar to mechanoadaptation since undifferentiated DMD myoblasts are indeed mechanically ill-equipped. Further, these results underscore a new influence of dystrophin in healthy myoblasts, even though we and others [64] find it is expressed at low levels (Figure 3 e). To our knowledge, human DMD myoblast traction forces have not been studied, yet, which underlines the potential of the new insights by quantifying the tensional homeostasis in the context of skeletal muscle diseases. Very recently, pathological DMD-related dystrophin mutations were reported to lead to lower focal adhesion tension in a FRET-based C2C12 model [65, 66]. Although this may sound contradictory to our results, it can be explained by larger focal adhesion sites, effectively distributing the increased forces over more molecules. Additionally, C2C12 cells with DMD-associated mutations have been reported to develop lower total strain energy, which also seems to be in conflict with our results. However, the total strain energy depends linearly on cell area, hence the reported slight decrease in strain energy might be a simple outcome of smaller C2C12-DMD cell size. Other investigations that measured the traction forces of iPSC derived cardiomyocytes did either only test the traction forces during a spontaneous contraction [67], or had only non-significant differences in Dystrophin expression [68]. Interestingly, Bremner et al. observe sarcomere shortening in dystrophic cardiomyocytes. We also identified a decreased sarcomere length in leg-derived bioengineered dystrophic skeletal muscle (Supplement 9) that correlates with increased homeostatic tissue tension which stresses the complexity of the emerging dystrophin-associated mechano-signaling.

### 2.4 Biomimetic DMD model exhibits pathological disease features

Throughout our study we found phenotypic variability and a large spread across our patient-derived data. Consistently, for myotube diameter (Figure 1 j), stimulated tissue contractility (Figure 2 f) and homeostatic tissue tension (Figure 2 h), we observe the most pronounced DMD phenotype when comparing leg muscle derived cell lines. By contrast, paravertebral derived bioengineered muscles did not exhibit expected DMD phenotypes to a similar extent. Whereas this may sound like a poor DMD model on the first glance, it in fact reveals the great sensitivity of it. Disease severity can depend on the muscle group being evaluated and DMD as a disease symptomatically starts in the lower extremities (e.g. quadriceps) and progresses to the upper body [9, 10, 12]. Additionally, it was shown that fast twitch skeletal muscle fibers are preferentially affected in DMD and first to degenerate in disease progression [69], which presents another explanation for our results. Connecting these findings to ours, we found larger portions of slow twitch Myosin heavy chain in our paravertebral derived bioengineered skeletal muscle (Supplement 1b) that showed less pronounced phenotypes. This leads us to the speculation that cells retain a memory of the muscle group they were derived from. Furthermore, we observed that in the paravertebral derived single myoblasts Utrophin levels were not decreased in the DMD background and *α*-7 Integrin isoform 3 was elevated, which suggest a compensatory mechanism (Supplement 7b,d). Paravertebral DMD5 cells even showed elevated levels of Utrophin and both isoforms of *α*-7 Integrin (Supplement 7b-d) which may account for the same mechanical features as its healthy control. All together, this may explain why we saw a more pronounced functional disability and deregulation of the mechanical homeostasis in leg derived bioengineered muscles throughout all of our experiments. To this end, our DMD model displays typical pathological features of the disease. In addition, the paravertebral muscles originate from an epaxial background, which are embryologically distinct from the hypaxial myotome (e.g. quadriceps) and may provide another reason for differences within our bioengineered muscles [70]. Thus, the muscular group origin of cell isolation seems to play a crucial role for bioengineering DMD models in the future. Since the immortalized patient-derived myoblasts originate from isolated muscle stem cells (MuSCs), we can speculate, that the MuSC population in dystrophic epaxial paravertebral muscles may not be as exhausted as in leg muscles.

### 2.5 Increased DMD homeostatic tissue tension is confirmed by isogenic models

Since patient derived cells exhibit immense phenotypical variability, we sought to demonstrate the close connection of the homeostatic tissue tension and functional strength to the dystrophin protein in three isogenic culture system. First, we investigated an established iPSC line from a Duchenne patient with an exon 48-50 deletion (’DMD51’) and compared it to the ‘Cor51’ iPSC line that was genetically corrected via exon 51 skipping (Figure 4 a) [48]. These isogenic cell lines were differentiated into skeletal myocytes by Shahriyari et al with a very similar myoblast/myocyte outcome for both cell lines [33]. We used cells from the exact same differentiation batch to bioengineer skeletal muscles in our glass bottom chamber over a period of 4 weeks.

**Fig. 4.**
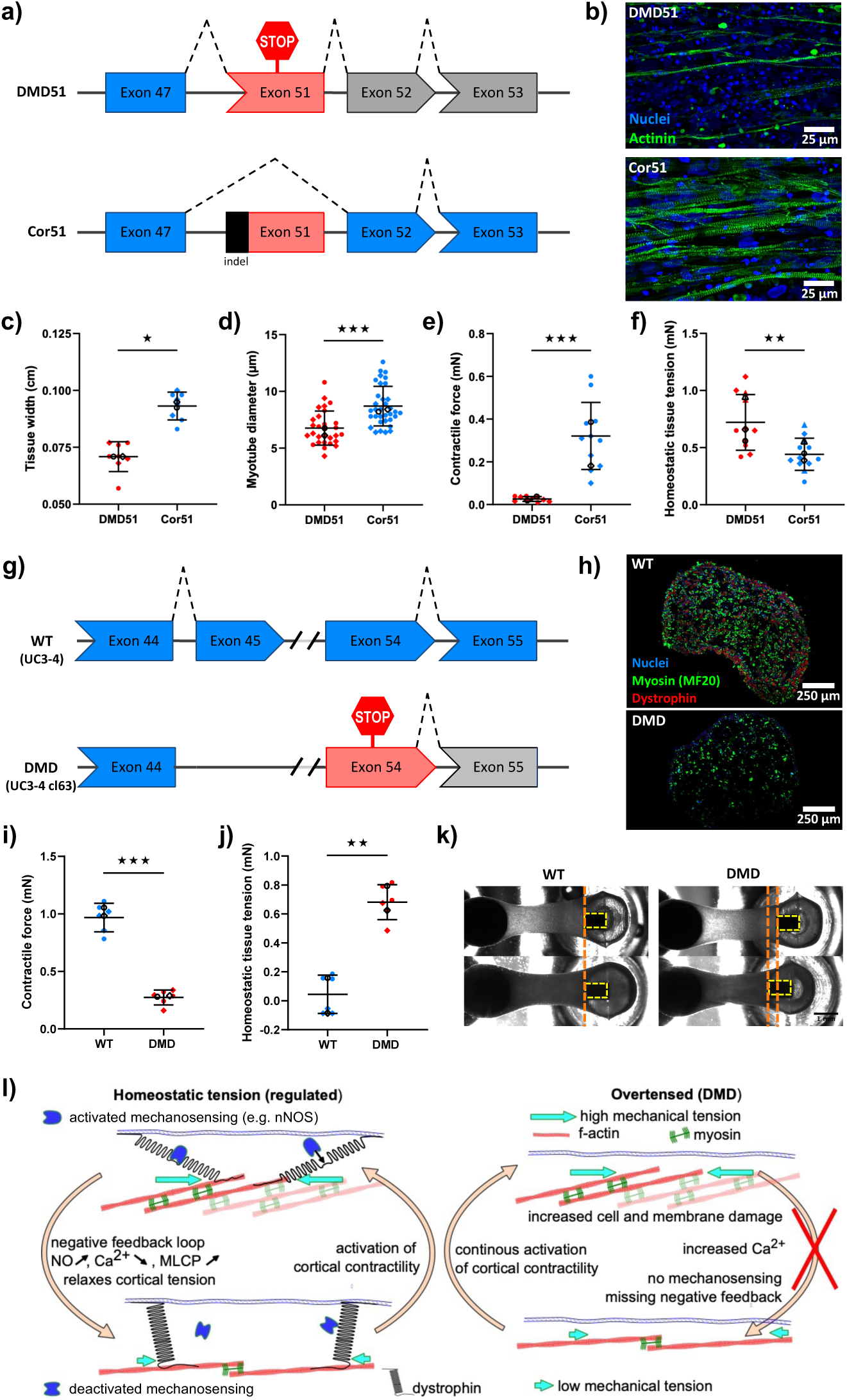
Exon 51 skipping in DMD51 restores tissue strength and tensional homeostasis. a) Schematic sketch of exon 51 skipping approach in iPSCs of DMD patients with exon 48-50 deletions. b) Representative flattened confocal stacks of multinucleated myotubes within DMD51 and Cor51 biomimetic muscles immunostained for sarcomeric-alpha-actinin (green) and nuclei counterstained with DRAQ7 (blue). c) Tissue width of DMD51 and Cor51 human biomimetic skeletal muscles. n=8 tissues per group from N=2 independent experiments d) Myotube diameter of DMD51 and Cor51 human biomimetic skeletal muscles. n=31/36 myotubes per group from N=2 tissue from two independent experiments each e) Contractile force of DMD51 and Cor51 human biomimetic skeletal muscles upon 20 Hz electrical stimulation. n=11/12 tissues per group from N=2 independent experiments f) Homeostatic tension of DMD51 and Cor51 human biomimetic skeletal muscles. n=11/12 tissues per group from N=3 independent experiments g) Schematic sketch of iPSC lines UC3-4cl63 (DMD) and healthy isogenic control UC3-4 (WT). h) Cross section images of WT and DMD microtissues generated in Curi Bio’s Mantarray platform and stained for myosin (green), dystrophin (red) and nuclei (blue). i) Contractile force of WT and DMD microtissue upon 100 Hz electrical stimulation. j) Homeostatic tension of WT and DMD human biomimetic skeletal muscle and associated representative bright field images (k). n=6 tissues per group from N=2 independent experiments l) Hypothetical molecular model of Dystrophin acting as a tension sensor that regulates the cells’ tensional homeostasis which is disturbed in DMD and leads to an increase in cellular tension mediated by cortical acto-myosin contraction. All results are presented as mean *±* SD. Statistical differences were analysed by unpaired t-test with p *<* 0.05 considered as significant. Independent experiments performed on different days are represented by different symbols, whereof the median of each independent experiment is presented in black.

Morphologically, we found that Cor51 showed multinucleated, striated myotubes in a more dense assembly when compared to DMD51 tissues (Figure 4 b). In addition, the myotubes formed from the Cor51 cells were also significantly thicker (Figure 4 d), which is in compliance with an overall higher macroscopic tissue width of the Cor51 engineered skeletal muscles (Figure 4 c). These dystrophin-dependent effects in the isogenic cell model are comparable to those we observed in patient derived pooled muscle tissues (Figure 1 g, i). This hints towards an elevated homeostatic tissue tension and a lower stimulated contractile strength of DMD51 bioengineered skeletal muscles when compared to the genetically corrected Cor51 tissues. As expected, the peak force of Cor51 engineered skeletal muscles was significantly higher following a 20 Hz electrical stimulation when compared to stimulated DMD51 tissues (Figure 4 e), supporting prior results on bioengineered heart muscle tissue showing that this dystrophin correction improves stimulated 3D engineered heart muscle tissue contraction [48]. Furthermore, we again find a significant increase in homeostatic tension in the DMD51 tissue, when compared to the Cor51 tissue (Figure 4 f). Similarly, we compared bioenginnered skeletal muscle tissues derived from the iPSC lines UC3-4 (’WT’) to its isogenic dystrophin-null (Figure 4 h) control UC3-4 cl63 (’DMD’) that was generated by deletion of Exon45 via CRISPR/Cas9, leading to a stop codon in Exon 54 (Figure 4 g)[71]. Unlike before, we bioengineered WT and DMD microtissues within a collagen-based matrix using Curi Bio’s Mantarray platform (Figure 4 k) in order to rigorously assess the independence of our findings from influences related to matrix composition and platform selection. Consistently, the contractile force of DMD tissue was significantly decreased (Figure 4 i) while the homeostatic tissue tension was significantly increased by even 7-fold (Figure 4 j). Interestingly single nuclei RNA sequencing of WT and DMD microtissue revealed differences in expression levels of calcium handeling (e.g RYR1) and integrin transcripts (e.g. ITGA4, ITGA5 or ITGBL1) after 12 days of culture (Supplement 10) suggesting both different mechanotransduction as well as perturbed intracellular calcium homeostasis, while myogenic markers were expressed in similar levels comparing 12 day old WT and DMD microtissue. Altogether, the results from both pair of isogenic iPSC-derived microtissue controls directly confirm our results in tensed dystrophic skeletal muscle tissues raised from patient-derived immortalized myoblasts.

Therefore, these results demonstrate that dystrophin, besides its shock absorbing function, plays a central role in the modulation of homeostatic tissue tension, and that this new, quantitative parameter should be included in any study of muscle function and malfunction. The increased homeostatic tension may also explain partially the higher cell death and membrane ruptures found in DMD muscles, as the cells are continuously operating at a much larger force generation, which might increase the probability of fiber damage. Indeed, these findings suggest a novel role for dystrophin as a tension regulator to establish an intact tensional homeostasis (Figure 4 l).

### 2.6 Dystrophin as a potential tension sensor

Myoblasts, myotubes and biomimetic skeletal muscles from a hypaxial dystrophic background exhibited elevated traction forces (Figure 3 b,c) and homeostatic tissue tension (Figure 2 h, Figure 4 f). Therefore, lack of a dystrophin protein displays a dysregulated tensional homeostasis in these cells, which suggests a tension sensor and regulator function for dystrophin (Figure 4 l). The high number of spectrin repeats that are known to unfold under critical pulling forces [15] suggests a direct mechanistic role of these, similar to the mechanosensing reported by the unfolding of similar spectrin repeats in *α*-actinin [16]. The tension-dependent unfolding and refolding capability of these spectrin-like repeats was recently demonstrated experimentally, and was interpreted as a shock absorbing function of dystrophin [72]. However, this exact mechanism may additionally reveal binding sites upon tension for downstream signaling proteins (Figure 4 l). To explain the observed downregulation of homeostatic tension in the presence of dystrophin, a molecule that is known to relax muscle contraction would be required to bind to a cryptic binding site that is only revealed if the tension surpasses a critical value. Indeed, dystrophin is known to interact with proteins that regulate mechanical loading in skeletal muscles and relaxation in smooth muscle (e.g. nNOS) [18, 45–47]. Hence, in line with this tension sensor model, it is not surprising that dystrophic cell lines show deregulated, and more specifically, elevated homeostatic tension. Our measurements support the hypothesis that dystrophin acts as a tension sensor where mechanical signaling proteins, like nNOS, may bind to dystrophin in a stretched or tensed condition and thus trigger the signaling that relaxes the muscle to maintain proper tensional homeostasis of healthy skeletal muscle (Figure 4 l). Since we showed that the homeostatic tension drastically decreases upon Blebbistatin and also CytochalasinD treatment (Figure 2 l,m), we believe that the major portion of the homeostatic tension is mediated by cortical acto-myosin contractility. This suggests that a lack of dystrophin in DMD and therefore misregulated mechanosensing and -signaling leads to a hypercontractile state of the cell cortex (Figure 4 l).

This new model provides a simple explanation for the surprising increase of homeostatic tension in dystrophic muscle, despite possessing reduced stimulated contractility when compared to healthy muscle. We suggest that a lack of dystrophin impairs intracellular sensing of critically high stresses. This deregulation may lead to an increased homeostatic tension which could result in elevated cellular damage. Assuming a limited force generation capacity of a muscle fiber, less contraction can be applied if the steady state, hence homeostatic, force generation is already elevated. This model is directly supported by the observed anticorrelation between homeostatic tension and stimulated force generation.

## 3 Conclusion

Here, we reported a novel mechanical key feature of Duchenne muscular dystrophy: Elevated homeostatic tissue tension (Figure 2 g, Figure 4 f,j). Our data suggest that tensional homeostasis within skeletal muscle tissue is important for proper functionality (Figure 2 i) since increased homeostatic tissue tension strongly correlates with decreased contractile strength upon stimulation. Furthermore, if the homeostatic, hence constant tension exceeds a maximal bearable level, this may furthermore explain the increased cell death as found in dystrophic muscles. Our results shine light on potential new fundamental functions of the dystrophin protein as a potential tension sensor protein (Figure 4 l) in addition to its proposed stabilizing function [72]. Moreover, our findings stress the importance of a highly regulated tensional homeostasis in skeletal muscle cells since we found that single dystrophic myoblasts show an increase in traction forces (Figure 3 c). In the future, follow up more clinically oriented studies might be based on this work to examine if the observed homeostatic tension increase is also relevant in humans. One can hypothesize that understanding the details of the potential mechanosensing by dystrophin might lead to new therapeutic targets aimed at interfering with muscle tension signaling to reduce characteristic progressive DMD muscle injuries and MuSC pool exhaustion as a strategy to prolong ambulation of patients.

## 4 Methods

### 4.1 PMMA chamber fabrication for 3D muscle tissue culture

Muscle tissue culture molds were fabricated from polymethyl methacrylate (PMMA) as previously described [26](ArtifiCell GmbH). In short, both the bottom part of the PMMA chamber containing 8 ellipsoid wells and the lid containing a pair of 16 mm long posts for each well were CNC milled. The bottom part was glued onto a microscopy cover glass (No1.5, VWR, Radnor, USA) using PDMS (Sylgard 170 or Sylgard 184 silicone, Sigma, St. Louis, USA). Prior to use, the molds were sterilized with 70 % ethanol and UV light and the wells were coated with a Poloxamere solution in ddH_2_O over night at 4 °C (5 % Pluronic F-127, Sigma, St. Louis, USA) to render the surface non-adhesive.

### 4.2 Human immortalized myoblast line cultivation

All immortalized human muscle progenitor cell lines (Table 1) were established at the Myoline platform of the Institut de Myologie (Paris, France) [49]. The cells were cultivated in tissue culture flasks (75 cm^2^, Greiner) in a Skeletal Muscle Cell Growth medium kit (PROMOCELL) supplemented with 15 % fetal calf serum (FCS, Sigma) and 1 % penicillin-streptomycin (Gibco) at 37 °C and 5 % CO_2_ in a humidified incubator. Skeletal Muscle Cell Growth medium was changed every other day. At 80 % confluency the cells were split using 0.25 % Trypsin-EDTA solution (Sigma).

### 4.3 Myogenic differentiation of iPSCs

Human induced pluripotent stem cell (iPSC) line DMD Del48-50 (DMD51) was purchased from RIKEN BioResource Center (HPS0164). The genetic correction of DMD51 by CRISPR/Cas9 mediated gene editing (Cor51) has been described before [48]. DMD51 and isogenic Cor51 lines were maintained on Matrigel-coated flasks in StemMACS iPSC Brew medium. Directed differentiation into skeletal myocytes was performed as previously reported by Shahriyari et al. 2022 [33]. In brief, iPSC were seeded in iPS-Brew XF medium with 5 *µ*mol/L of Y27632 (Stemgent) for 24 hrs to obtain a 30 % confluent culture. To start differentiation, medium was switched to DMEM with 1 % Pen/Strep, 1 % N-2 Supplement, 1 % MEM non-essential amino acid solution (N2 basal medium, all Thermo Fisher Scientific), 10 *µ*mol/L CHIR99021 (Stemgent), 0.5 *µ*mol/L LDN193189 (Stemgent), and 10 ng/mL FGF-2 (Peprotech) for 4 days. At day 4, the medium was exchanged with N2 basal medium, 20 ng/mL FGF-2, and 10 *µ*mol/L DAPT (Tocris) until day 6. On day 6 and 7, 10 ng/mL HGF (Peprotech) was added to the medium. The medium was then switched to N2 basal medium, 10 *µ*mol/L DAPT, 10 ng/mL HGF, and 10 % knockout serum replacement (Thermo Fisher Scientific) on days 8, 9, 10 and 11. From day 13 to 22, myogenic cells were expanded in N2 basal medium, 10 % knockout serum replacement, and 10 ng/ml HGF. On Day 22, skeletal myocytes were then enzymatically dissociated with TrypLE (Thermo Fisher Scientific) for 5 to 7 minutes at 37 °C and frozen in expansion medium with 5 *µ*mol/L Y27632 (Stemgent). Similarly, skeletal muscle myoblasts were generated from the UC3-4 DMD null and isogenic control iPSC lines as previously reported [71, 73].To initiate differentiation, iPSCs were seeded at 15,000 cell-s/cm² in mTeSR Plus and grown to 40 % confluency. On day 0, the culture medium was replaced with a differentiation medium consisting of DMEM/F12 (Gibco No:10565018), 1x non-essential amino acids (Gibco No:11140050), 1x Insulin-Transferrin-Selenium (Gibco No:41400045), 3 *µ*M CHIR99021 (Axon No:1386), and 0.2 *µ*g/mL LDN193189 (Miltenyi Biotec No:130-106-540). To support progenitor expansion, 20 ng/mL bFGF (R&D Systems No:3718-FB) was added to the culture on day 2. The transition to myoblast maturation began on day 5 with a medium change, where CHIR99021 and ITS were withdrawn and replaced by 15 % KSR (Gibco No:10828028), 2 ng/mL IGF-1 (R&D Systems No:291-G1), and 10 ng/mL HGF (R&D Systems No:294-HGN). The maturation medium was finalized on day 7 with the removal of both bFGF and LDN193189. Cells were maintained in this final medium until day 28, with feeds every other day. For expansion, the confluent sheet was mechanically dissociated, replated onto a Matrigel-coated T225 flask, and cultured in skeletal muscle growth media (SKGM). On day 32, cells were purified by fluorescence-activated cell sorting (FACS) for the NGFR-positive population (Biolegend No:345108) after being passed through a 40*µ*m filter. Experiments were performed using cells between passages 4 and 5.

### 4.4 3D human skeletal muscle microtissue cultivation

3D skeletal muscle tissues were raised in culture as previously reported by various groups [23, 24, 26]. Briefly, 1.5 × 10^7^ cells per mL were resuspended in an ECM mixture consisting of DMEM (40 % v/v, Capricorn, Ebsdorfergrund, Germany), 4 mg/mL bovine fibrinogen (Sigma) in 0.9 % (w/v) NaCl solution in water and Geltrex™ (20 % v/v, Gibco). 25 *µ*L of the cell mixture was used to seed each tissue. Fibrin polymerisation was induced with 0.5 units thrombin (Sigma) per mg of fibrinogen for 5 min at 37 °C. Subsequently, 400 *µ*L Skeletal Muscle Cell Growth medium supplemented with 1.5 mg/mL 6-aminocaproic acid (ACA, Sigma) was added. After 2 days, the growth medium was exchanged to differentiation medium containing DMEM supplemented with 2 % horse serum (HS, Sigma), 1 % penicillin-streptomycin (Gibco), 2 mg/mL ACA and 10 *µ*g/ml human recombinant insulin (SAFC Biosciences). The differentiation medium was changed every other day. DMD51 and Cor51 derived bioengineered skeletal muscles were reconstituted in expansion medium (N2 basal medium, 10 % knockout serum replacement, and 10 ng/ml HGF) for one week followed by maturation in DMEM with 1 % Pen/Strep, 1 % N-2 Supplement, and 2 % B-27 Supplement (maturation medium) for additional three weeks. UC3-4 (WT) and isogenic UC3-4cl63 (DMD) derived microtissue were generated in the Mantarray platform (Curi Bio, USA) in a 1.3 mg/mL rat tail collagen I based matrix adapted from Smith et al [71]. In short, iPSC-derived myoblasts were combined with human dermal fibroblasts (Lonza, CC-2511) at a 9:1 ratio. Following centrifugation (300 × g, 5 min), cells were resuspended to achieve 420,000 cells/construct. Cell suspensions were incorporated into a hydrogel matrix containing rat tail collagen I (1.3 mg/mL; Sigma-Aldrich, 08-115), Matrigel (20 %; Corning, 356231), MEM (10 %; Gibco, 11095080), and DMEM with 4.5 g/L glucose plus Glutamax (10 %). Sixty microliters of this mixture was dispensed into Mantarray casting trenches. After polymerization (30 min, 37°C, 5 % *CO*_2_), constructs were cultured in SKGM for 24 h. Tissues were subsequently transferred to differentiation conditions using SKDM containing DMEM/Glutamax, Knockout Serum Replacement (2 %; Gibco, 10828028), B27 supplement (1×; Gibco, 17504001), IGF-1 (10 ng/mL; R&D Systems, 291-G1), and HGF (10 ng/mL; R&D Systems, 294-HGN). Media exchanges occurred every 48-72 h for 4 weeks.

### 4.5 Immunostaining and confocal fluorescence microscopy

Confocal microscopy and immunohistological investigations were performed as previously reported [26]. In brief, human 3D skeletal muscle tissues were fixed using 4 % paraformaldehyde (PFA) for 15 min at room temperature within the culture mold. Next, the samples were blocked for 1 h at room temperature using PBS supplemented with 20 % goat serum (GS, Sigma) and 0.2 % Triton-X-100 (Carl Roth). Afterwards, the tissues were incubated with the primary antibody (monoclonal mouse anti sarcomeric alpha actinin, 1:100, Abcam; monoclonal mouse anti Dystrophin, 1:100, Santa Cruz Biotechnology, monoclonal mouse anti fast myosin skeletal heavy chain, Abcam; monoclonal mouse anti slow myosin skeletal heavy chain, Abcam) diluted in blocking solution over night at 4 °C. Subsequently, the tissues were washed with blocking solution three times and incubated with the respective secondary antibody (polyclonal goat anti-mouse IgG1, 1:500, Invitrogen) for 45 min at room temperature. Cell nuclei were counterstained using DRAQ7 (1:100, Abcam) or Hoechst33342 (Thermo Fisher Scientific). Confocal images were acquired using Slidebook 6 software (3i) using an inverted microscope (Nikon Eclipse Ti-E) equipped with a CSU-W1 spinning disk head (Yokogawa) and a scientific CMOS camera (Prime BSI, Photometrics). Images were analysed and prepared for publication using the open source software Fiji [74].

### 4.6 Sarcomere lengths analysis

For the analysis of sarcomere lengths, we utilized deep-learning-based detection of Z-bands in conjunction with double-wavelet analysis, employing the SarcAsM package [75].

### 4.7 Single nuclei RNA sequencing

Single nuclei were released from bioengineered skeletal muscle microtissues or myogenic progenitors derived from iPSC after incubation in collagenase and dispase (2 hours at 37°C), accutase (1h at 37°C), cell dissociation and incubation in lysis buffer (Tris-HCl, NaCl, MgCl2, Tween20 0.1 %, Igepal 0.1 %, Digitonin 0.01 %, BSA 1 %, RNase Inhibitor). The single nuclei RNA-Seq library was then prepared by split-pool barcoding using the Parse Biosciences technology and kits. Data were demultiplexed and aligned using the Parse Biosciences pipeline, and then analyzed with the Seurat suite (v5.3.0).

### 4.8 Electrical stimulation of biomimetic human skeletal muscle tissues

Contractile function of human 3D skeletal muscle tissues was examined two weeks after differentiation of immortalized myoblasts or four weeks after iPSC-derived tissue maturation. For that purpose, a metal pin (0.55 mm in diameter, Fine Science Tools) was implanted behind each post as electrodes. Via copper wires the electrodes were directly connected to a custom built signal generator accessed by a custom written program [76]. The tissues were stimulated using square-wave pulses with 20 % duty cycle, 5 V amplitude and 20 Hz frequency (tetanus contraction). Time lapse videos of the posts during contraction were subsequently used for post deflection analysis. Contractile function of UC3-4 (WT) and isogenic UC3-4cl63 (DMD) tissue was measured using the Mantarray platform (Curi Bio) 3 times a week. EMT plates (24-well) were placed in the instrument’s heated measurement chamber at 37°C and fitted with a stimulation lid that positioned electrodes in each well for simultaneous electrical pacing (Tetanic contractions at 100Hz, 5 ms pulse width, 100 mA). The Magnetic field flux was sampled at 100 Hz and converted to post deflection (mm) via the manufacturer’s magnet localization algorithm, from which contractile force and force-time traces were derived. T. Raw force data was analyzed using the Engineered Muscle Contractility Analysis Pipeline (EMCAP v1.0), a custom open-source Python-based peak-finding and analysis package [77].

### 4.9 Homeostatic tension analysis of biomimetic human skeletal muscle tissues

The homeostatic tension of human 3D skeletal muscle tissues was evaluated two weeks after differentiation of immortalized myoblasts or four weeks after iPSC-derived tissue maturation. For that purpose, the distance of the two posts was measured directly after the seeding procedure and after tissue maturation. The reduction in post distance was multiplied by the posts’ spring constant (39 *µN/µm*, [26]) to obtain the homeostatic tissue tension. The distances of the posts was measured using custom written programs [78, 79](ArtifiCell GmbH). The time and concentration dependent release of homeostatic tissue tension upon drug addition was analysed using the custom program [79]. In the rare event of a negative tension outcome the measurement was discarded.

### 4.10 Post deflection analysis

For post deflection analysis, we proceeded as previously described [26]. In brief, we focused on the edge of the post with our imaging system during tissue contraction and recorded a time series. Following the deflection of the post throughout the whole time series with custom written programs [80, 81](ArtifiCell GmbH), the pulling forces were determined by multiplication of the post displacement by the apparent spring constant of the post (39 *µN/µm*, [26]).

### 4.11 Length-tension relationship analysis

To generate ring-shaped bioengineered skeletal muscle, immortalized myoblast cell lines AB1167, KM571, and AB1023 were cast in a 250 *µ*l ring-format mold [33] utilizing a fibrin/geltrex hydrogel as described above. After hydrogel compaction for 7 days inside the mold, the bioengineered muscles were transferred onto static stretchers for mechanical loading and further maintained for 14 days in differentiation medium. To evaluate the Young’s modulus, the tissues were subjected to step-wise length increases in thermostatted organ baths filled with Tyrode’s solution. The baseline stress for each tissue was set individually to 0.2 mN. The Young’s modulus (E) was calculated by measuring the baseline force (F) under a given deformation (d) and determining the cross-sectional area (A) of the tissue with L being the length of the tissue: *E* = *F/A* ∗ *L/d*

### 4.12 Traction force microscopy (TFM)

In this study, the traction forces of healthy and DMD myotubes and single human myoblasts were examined as previously described [82]. For differentiated myotubes we used 36 kPa and for single myoblasts 15 kPa (Young’s modulus) polyacrylamide (PAA) gels with 100 nm polystyrene beads (1:50, micromod) embedded. Briefly, TFM of single myoblasts was performed 12-18 hours after seeding 1 × 10^5^ cells on the fibronectin (Sigma) coated PAA gel surface within a 35 mm glass bottom dish (Greiner). For myotube TFM, 8 × 10^5^ cells were seeded in Skeletal Muscle Cell Growth medium and the next day differentiated for one week in differentiation media. PAA gels and covalent ECM coating was achieved similarly to the methods described previously [82]. Unless stated otherwise, all chemicals were obtained from Sigma-Aldrich (Steinheim, Germany). Polyacrylamide (PAA) gels were prepared following the method described by Bollmann et al. (2015), with some modifications. Glass bottom dishes (CELLview 35/10 mm, Greiner Bio-One International, Kremsmünster, Austria) were first cleaned with 70% ethanol, followed by 0.1 N NaOH. The glass bottom was then coated with 200 *µL* of (3-Aminopropyl) trimethoxysilane (APTMS) for 3 minutes, thoroughly rinsed, and covered with 500 *µL* of 0.5% glutaraldehyde for 30 minutes. Meanwhile, the PAA gel premix was prepared by adding 4 *µL* of acrylic acid to a mixture of 250 *µL* of 2% N,N’-methylenebisacrylamide and 500 *µL* of 40% acrylamide solution. For 15 (30) kPa stiff PAA gels, 113 (150) *µL* of this mixture were gently combined with 377 (340) *µL*of 65% PBS and 10 *µL* of a fluorescent bead solution (100 nm NH2-coated micromer-redF, Micro-mod, Rostock, Germany). Polymerization was initiated by adding 5 *µL* of 10% ammonium persulfate solution (APS) (Roth, Karlsruhe, Germany) and 1.5 *µL* of N,N,N’,N’-tetramethylethylenediamine (TEMED) (Roth, Karlsruhe, Germany). To functionalize the gels for coating, the acrylic acid was activated with a solution of 0.2 M N-(3-dimethylaminopropyl)-N-ethylcarbodiimide hydrochloride (EDC), 0.1 M N-hydroxysuccinimide (NHS), 0.1 M 2-(N-Morpholino) ethanesulfonic acid (MES), and 0.5 M NaCl for 15 minutes at room temperature, followed by thorough washing with PBS and incubation with fibronectin (50 *µ*g/mL) at 37 °C for 1 hour or at 4 °C overnight. The gel’s stiffness was verified by rheological measurements (AFM and Rheometer) to be consistent with the protocol.

Utilizing a 40X WI objective two z-stack images were acquired, one showing the cells on the substrate, the second image after cell lysis induced by addition of 500 *µL* of 5 % SDS. The bead displacement between the two images was used to calculate forces and deformation energies using Elastix software [83, 84] embedded in a custom-made MATLAB (MathWorks) script [62].

To obtain the traction forces, an adapted version of the Fourier Transformation based traction force microscopy was used, where the finite thickness of the PAA gel was accounted for using the method introduced by [85], and the Tikhonov’s regularization parameter was optimized using the background noise in an area far away from the cells using the Bayesian inference approach by Huang et al [63]. The evaluation of traction forces and the resulting strain energy and contractile moments were restricted to the cell-area set by previously defined masks which cover the cell area. The total strain energy was calculated by integrating the scalar product of traction stress and deformation over the cell area. The normalized strain energy divided the total strain energy by the area of the cell. The contractile moment was obtained following the approach by Butler et al [59]. Briefly, the shear moment matrix was determined by integrating the product of the vectoral components of position and force: *M_ij_* = 1*/*2 ∫ *dr*^2^(*x_i_T_j_*(*r*) + *x_j_T_i_*(*r*)) and then using the trace of this matrix as these are the contractile elements.

### 4.13 Mass spectrometry-based proteome profiling

Protein samples were prepared by lysis of 5 × 10^6^ cells/mL for 5 min at 95 °C in SDS sample buffer composed of 62.5 mM Tris/HCl (pH 6.8), 10 % glycerol, 2 % sodium dodecyl sulfate, 5 % beta-mercapto-ethanol and 0.005 % bromphenol blue. The samples were loaded onto a 4-12 % NuPAGE Novex Bis-Tris Minigels (Invitrogen) and run into the gel for 1.5 cm. Following Coomassie staining, the protein areas were cut out, diced and subjected to reduction with dithiothreitol, alkylation with iodoacetamide and finally overnight digestion with trypsin. Tryptic peptides were extracted from the gel, the solution dried in a Speedvac and kept at −20°C for further analysis c. Protein digests were analyzed on a nanoflow chromatography system (nanoElute) hyphenated to a hybrid timed ion mobility-quadrupole-time of flight mass spectrometer (timsTOF Pro, all Bruker). In brief, 400 ng equivalents of peptides were dissolved in loading buffer (2 % acetonitrile, 0.1 % trifluoroacetic acid in water), enriched on a reversed-phase C18 trapping column (0.3 cm × 300 *µ*m, Thermo Fisher Scientific) and separated on a reversed-phase C18 column with an integrated CaptiveSpray Emitter (Aurora 25 cm × 75 *µ*m, IonOpticks) using a 100 min linear gradient of 5-35 % acetonitrile / 0.1 % formic acid (v:v) at 250 nL/min, and a column temperature of 50◦C. Both identification and quantification was achieve by directDIA analysis in diaPASEF mode [86] using 20 variable width isolation windows from m/z 400 to 1,350 to include the 2+/3+/4+ population in the m/z–ion mobility plane. The collision energy was ramped linearly as a function of the mobility from 59 eV at 1/K0=1.5 Vs cm*^−^*^2^ to 20 eV at 1/K0=0.7 Vs cm*^−^*^2^. Two technical replicates per biological replicate were acquired. Protein identification was achieved using the Pulsar algorithm in Spectronaut Software version 16.0 (Biognosys) using default settings. All DIA data were searched against the UniProtKB Homo sapiens reference proteome (revision 01-2021) augmented with a set of 51 known common laboratory contaminants at default settings. For quantitation, up to the 6 most abundant fragment ion traces per peptide, and up to the 10 most abundant peptides per protein were integrated and summed up to provide protein area values. Mass and retention time calibration as well as the corresponding extraction tolerances were dynamically determined. Both identification and quantification results were trimmed to a False Discovery Rate of 1 % using a forward-and-reverse decoy database strategy.

### 4.14 Mouse Ethics and Studies

Animal use protocols were reviewed and approved by the local Animal Care Committee (ACC) within the Division of Comparative Medicine (DCM) at the University of Toronto. All methods in this study were conducted as described in the approved animal use protocol (#20012838) and more broadly in accordance with the guidelines and regulations of the DCM ACC and the Canadian Council on Animal Care. Mice were housed with a 14-hour light/10-hour dark cycle and were provided with ad libitum access to water and food. C57Bl/10ScSn-DMD^mdx/J and C57Bl/10ScSn mice were purchased from The Jackson Laboratory (USA) and utilized in experiments after a 1-week acclimatization period.

### 4.15 Statistical analysis

Results are presented as mean ± SD (exceptions: Figure 2 i, Figure 3 d, Supplement 4 & Supplement 8b, mean ± SEM) and statistical differences of experimental groups were analysed by one way ANOVA multiple comparison or unpaired two-sided t-test using GraphPad Prism software. All data were analysed regarding normal distribution with GraphPad Prism using the Shapiro-Wilk test. For not normally distributed data the Kruskal-Wallis or Mann-Whitney test was used, respectively. p *<* 0.05 was considered as significant, and significances were subdivided into three levels: ★ (p = 0.05 - 0.01), ★★ (p = 0.01 - 0.001), ★★★ (p *<* 0.001). Pearson correlation coefficient was calculated using GraphPad Prism. The number of replicates (n) and independent experiments (N) is indicated in every figure legend.

## 5 Declarations

ADH, ML,TMM and TB have recently founded a company, ArtifiCell GmbH, which provided some of the equipment used in this study. The authors declare that this relationship does not influence the objectivity or integrity of the research presented. A patent application has been filed for the chamber: WO/2021/229097, PCT/EP2021/062976. The other authors declare no conflict of interest.

## Acknowledgments

The authors thank Vincent Mouly and the Myoline platform from the Institut de Myologie, Paris for generating the cells used. Additionally, we thank the Proteomics Core Facility of the University Medical Center in Göttingen for their help with mass spectrometry analysis. This research was conducted within the Max Planck School Matter to Life (via PM) supported by the German Federal Ministry of Education and Research (BMBF) in collaboration with the Max Planck Society. This work was funded by the Human Frontiers Science Program (to PMG and TB). PMG is the Canada Research Chair in Endogenous Repair. Funding to PMG is from the Natural Sciences and Engineering Research Council and Medicine by Design, a Canada First Research Excellence Program. TB was supported by the European Research Council (consolidator grant number 771201) and the Deutsche Forschungsgemeinschaft (DFG) under the project numbers 450595133; 456112451. We acknowledge support by the Open Access Publication Funds of the Göttingen University. Mass spectrometry equipment used in this study was jointly funded by the Deutsche Forschungsgemeinschaft (DFG) and the State of Lower Saxony under the project number 442069358.

## Data and code availability

All supporting data are available from the authors upon request. Custom code used in this study is available on github: https://github.com/Tillmuen09/, https://github.com/mazelbr/.

## 7 Supplementary information

Supporting information is available from the authors upon request.

**Supplement 1:**
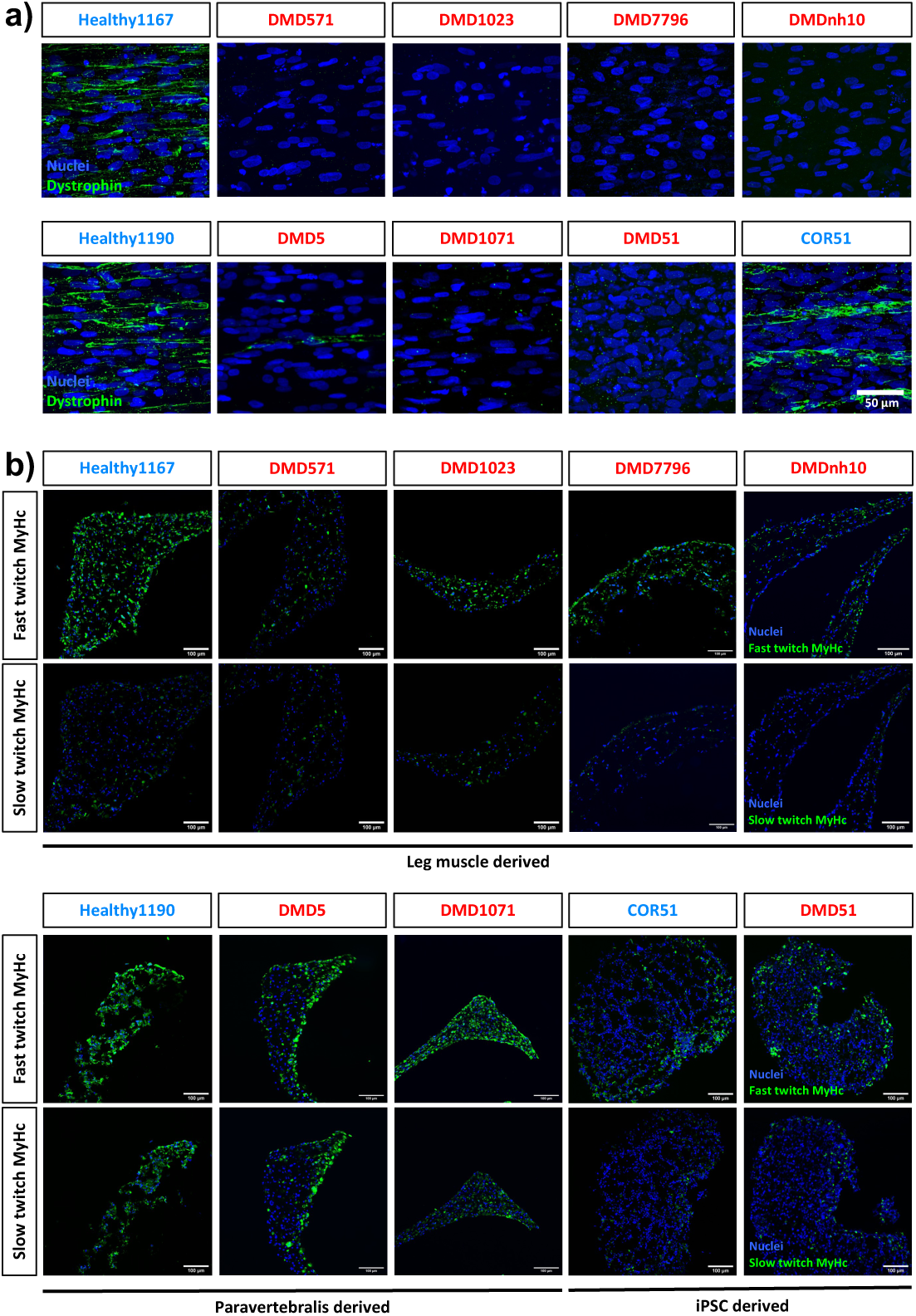
a) Representative flattened confocal stacks of healthy and DMD human bioengineered skeletal muscle tissues immunostained for dystrophin (green) and nuclei counterstained with DRAQ7 (blue). b) Representative confocal microscopy cryosection image of healthy and DMD human bioengineered skeletal muscle tissues immunostained for fast and slow twitch myosin heavy chain (green) and nuclei counterstained with Hoechst33342 (blue).

**Supplement 2:** Video of a 20 Hz tetanus contraction of a Healthy1167 bioengineered skeletal muscle tissue two weeks after differentiation.

**Supplement 3:** Video of a post deflection upon a 20 Hz tetanus contraction of a Healthy1167 bioengineered skeletal muscle tissue two weeks after differentiation.

**Supplement 4:**
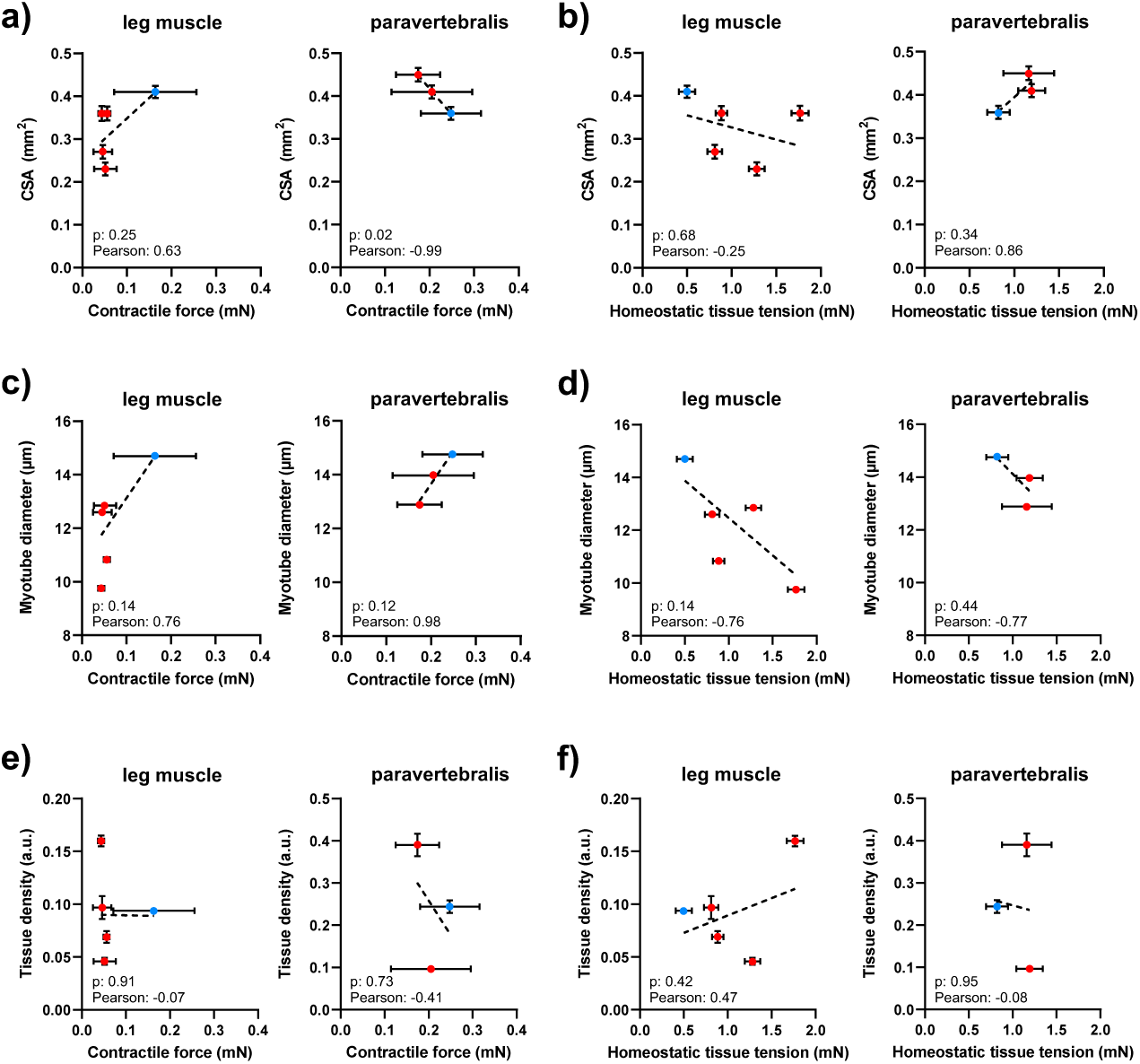
a) Correlation between the cross sectional area and the contractile force upon a 20 Hz electrical stimulation for leg and paravertebralis derived microtissues. b) Correlation between the cross sectional area and the homeostatic tissue tension for leg and paravertebralis derived microtissues. c) Correlation between the myotube diameter and the contractile force upon a 20 Hz electrical stimulation for leg and paravertebralis derived microtissues. d) Correlation between the myotube diameter and the homeostatic tissue tension for leg and paravertebralis derived microtissues. e) Correlation between the tissue density and the contractile force upon a 20 Hz electrical stimulation for leg and paravertebralis derived microtissues. f) Correlation between the tissue density and the homeostatic tissue tension for leg and paravertebralis derived microtissues. All results are presented as mean ± SEM.

**Supplement 5:**
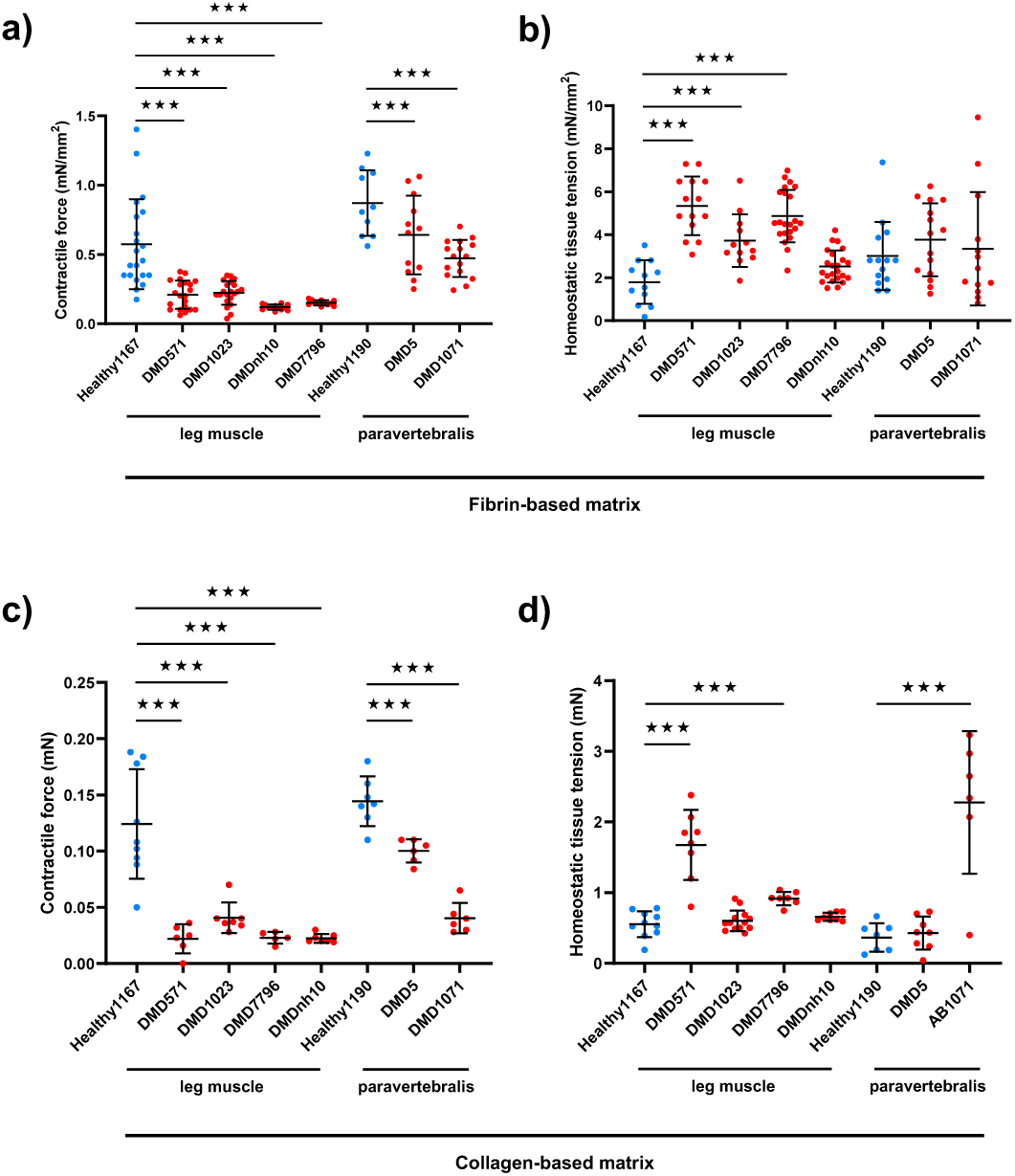
a) Specific contractile force normalized on entire tissue CSA of healthy and DMD human skeletal muscle tissues raised in fibrin-based matrix upon 20 Hz electrical stimulation, separated between the different patients (n=21/21/21/13/12/10/12/16 tissues per group from N=3 independent experiments each). b) Specific homeostatic tissue tension normalized on entire tissue CSA of healthy and DMD human bioengineered skeletal muscle tissues raised in fibrin-based matrix, separated between the different patients (n=12/14/12/22/23/14/15/13 tissues per group from N=3 independent experiments each). c) Contractile force of healthy and DMD human skeletal muscle tissues raised in collagen-based matrix upon 20 Hz electrical stimulation, separated between the different patients (n=9/6/7/5/7/6/6 tissues per group from N=2 independent experiments each). d) Homeostatic tissue tension of healthy and DMD human skeletal muscle tissues raised in collagen-based matrix, separated between the different patients (n=10/8/13/7/9/8/6 tissues per group from N=2 independent experiments each). Statistical differences were analysed by one way ANOVA multiple comparison with p *<* 0.05 considered as significant. All results are presented as mean ± SD.

**Supplement 6:**
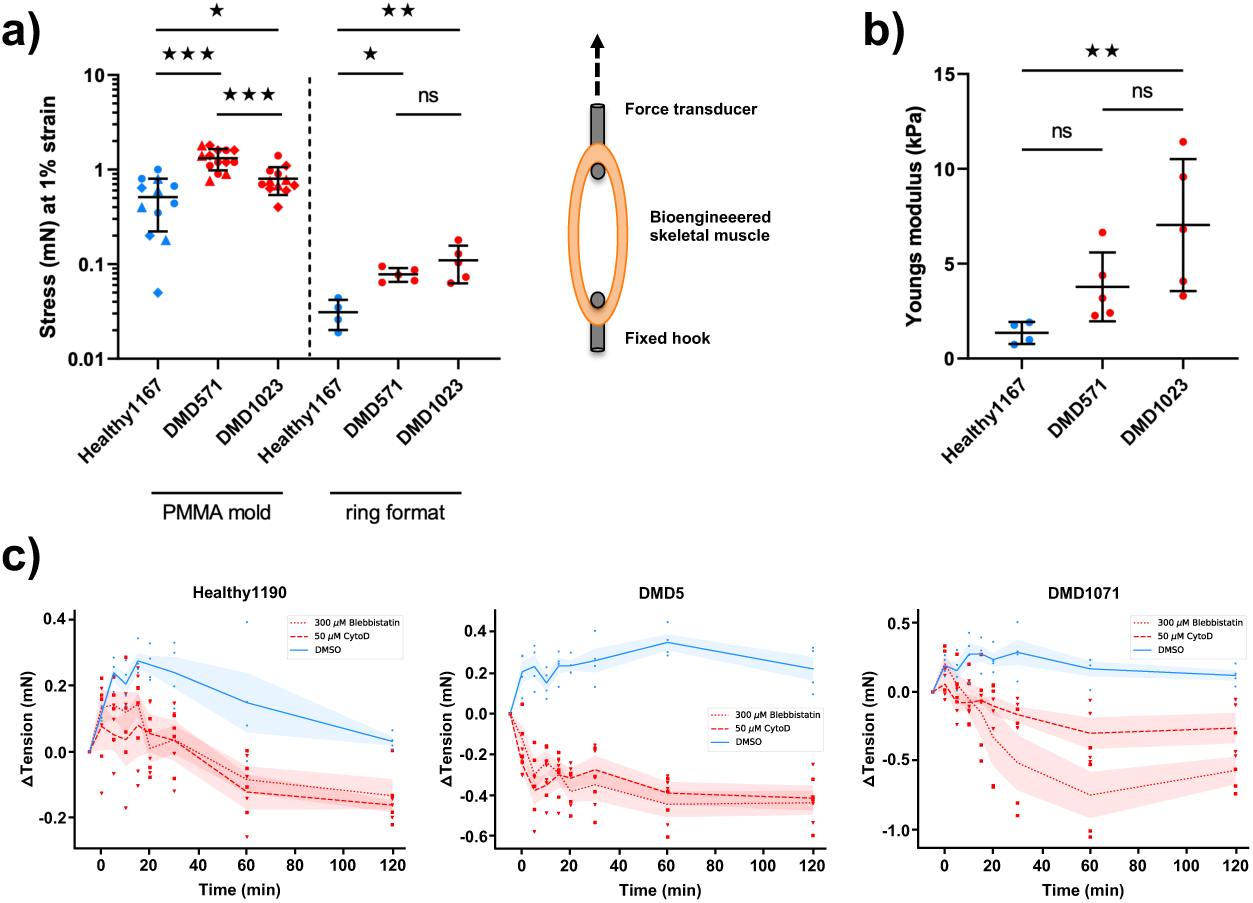
a) Length-tension relationship comparison of bioengineered skeletal muscle tissues in our PMMA mold (n=12/14/12 tissues per group from N=3 independent experiments each) and a ring format (n=4/5/5 tissues per group from N=1 single experiment each). b) Young’s modulus of the ring shaped bioengineered skeletal muscles obtained from organ bath length-tension relationship measurements (n=4/5/5 tissues per group from N=1 single experiment each). c) Homeostatic tissue tension decrease upon 300 *µ*M Blebbistatin and 50 *µ*M CytochalasinD treatment (red) compared to DMSO control tissues (blue) for paravertebralis derived myoblast lines Healthy1190, DMD5 and DMD1071 (n=4 tissues per condition from N=1 independent experiments). Results in c) are presented as mean ± SEM, all other results as mean ± SD. Statistical differences were analysed by one way ANOVA multiple comparison with p *<* 0.05 considered as significant.

**Supplement 7:**
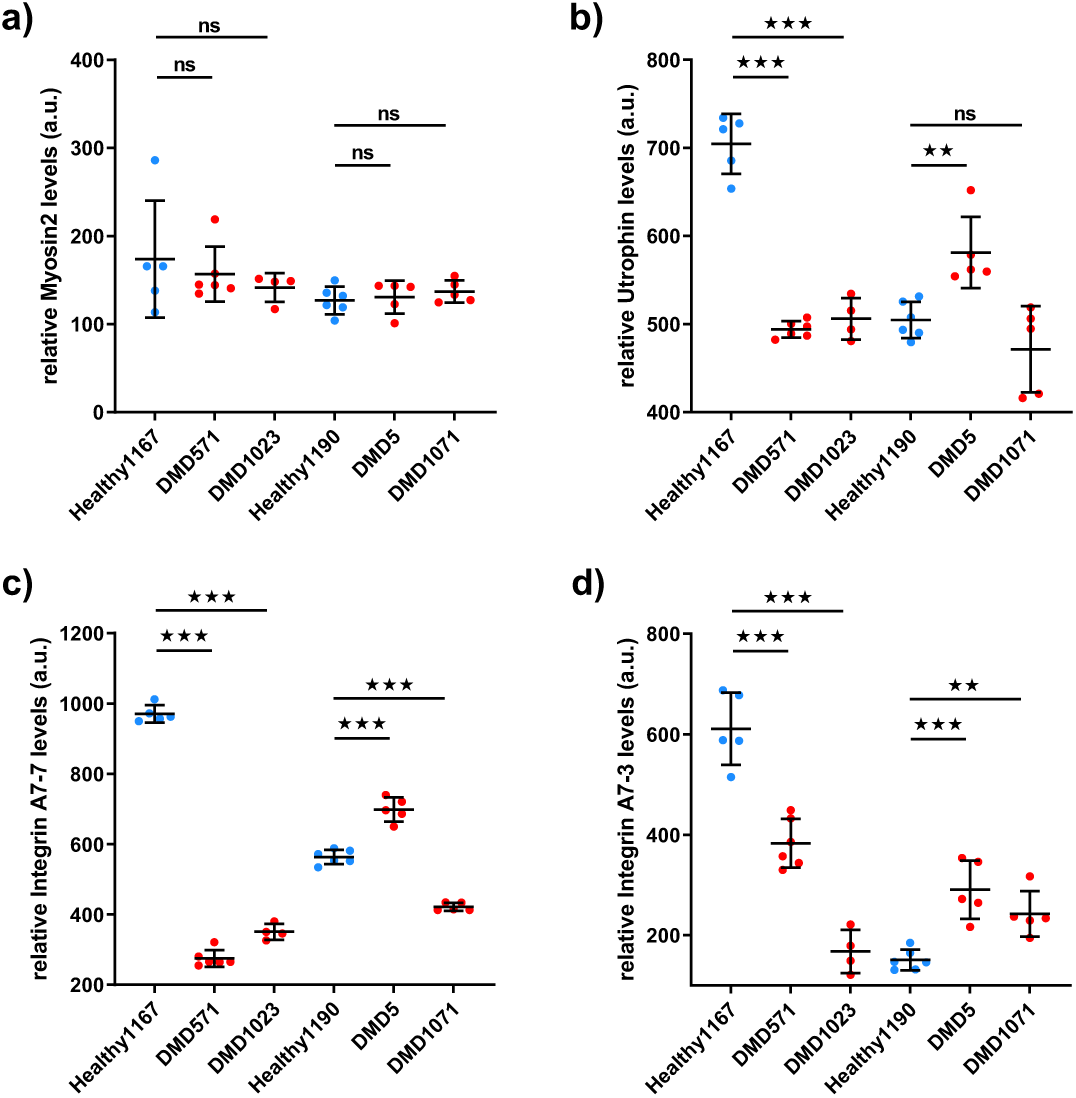
Changes in protein levels of healthy and DMD myoblasts determined by data independent acquisition mass spectrometry (DIA-MS) for Myosin2 (a), Utrophin (b), *α*-7 Integrin isoform 7 (c) and 3 (d) (n=5/6/4 samples per group collected from N=1 independent experiment). Statistical differences were analysed by one way ANOVA multiple comparison with p *<* 0.05 considered as significant. All results are presented as mean ± SD.

**Supplement 8:**
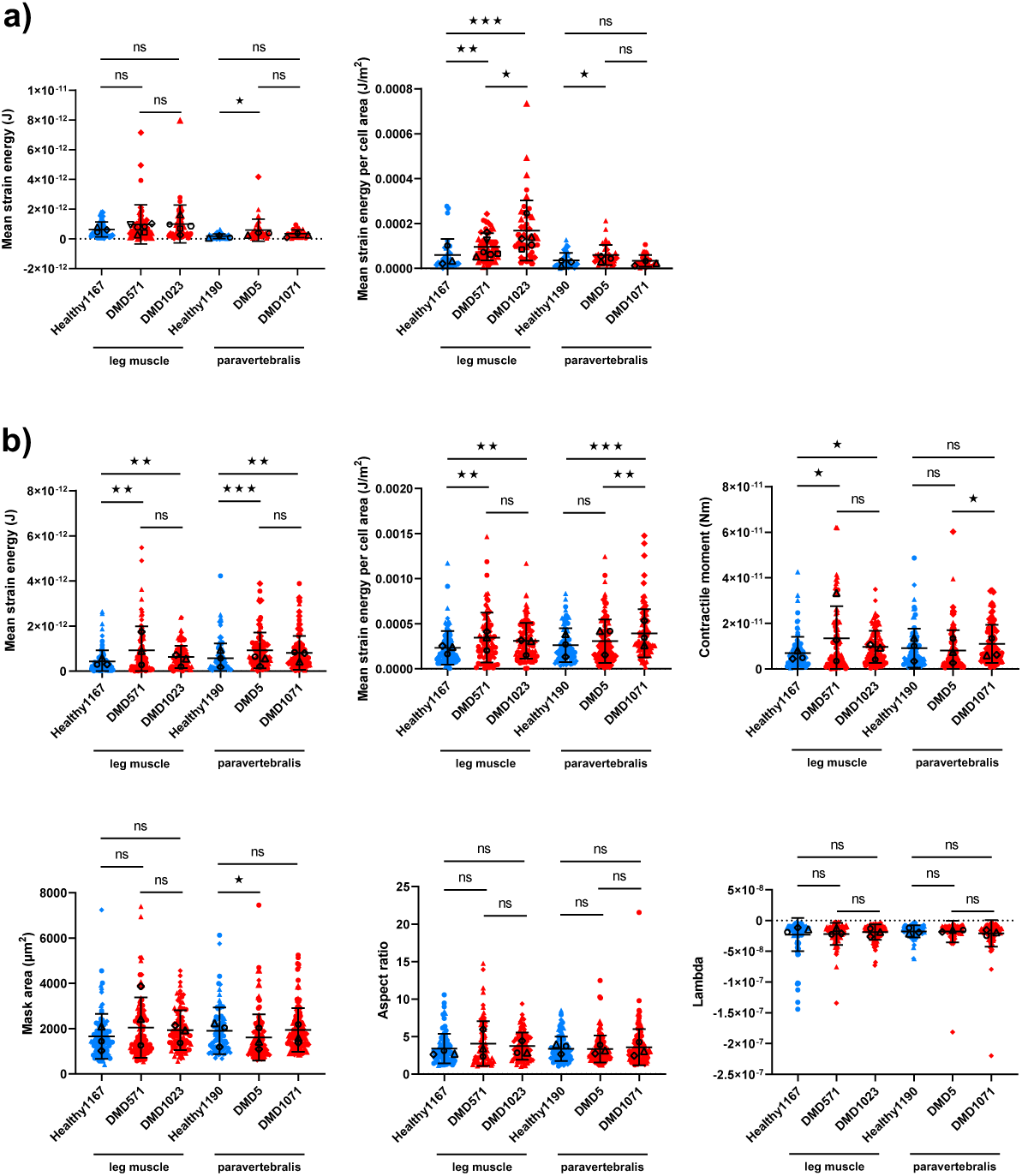
a) Mean strain energy and mean strain energy per cell area of healthy and DMD myotubes (n=34/46/46/38/38/33 myotubes per group from N=3/6/5/3/3/3 independent experiments). b) Mean strain energy, mean strain energy per cell area, contractile moment, mask area, aspect ratio and regularisation parameter “Lambda” of healthy and DMD single myoblasts TFM (n=94/81/108/89/103/119 myoblasts per group from N=3 independent experiments each). Statistical differences were analysed by one way ANOVA multiple comparison with p *<* 0.05 considered as significant. Independent experiments performed on different days are represented by different symbols, whereof the median of each independent experiment is presented in black. All results are presented as mean ± SD.

**Supplement 9:**
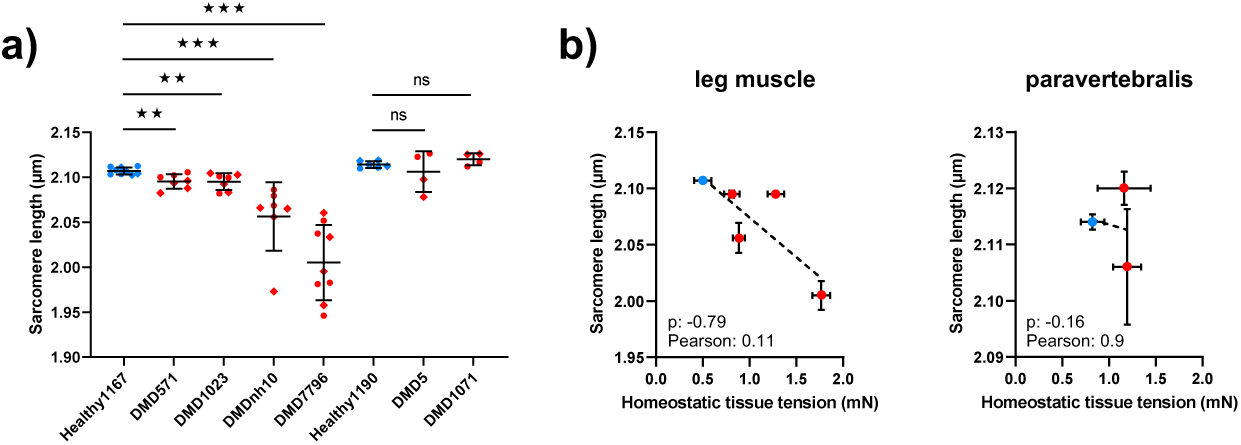
a) Sarcomere length within healthy and dystrophic bioengineered skeletal muscle (n=9/7/7/7/9/6/4/4 confocal microscopic z-stack acquisitions per group from N=2 tissue from two independent experiments each). b) Correlation between the Sarcomere length and the homeostatic tissue tension for leg and paravertebralis derived microtissue. Statistical differences were analysed by one way ANOVA multiple comparison with p *<* 0.05 considered as significant. Independent experiments performed on different days are represented by different symbols. Results in a) are presented as mean ± SD and in b) as mean ± SEM.

**Supplement 10:**
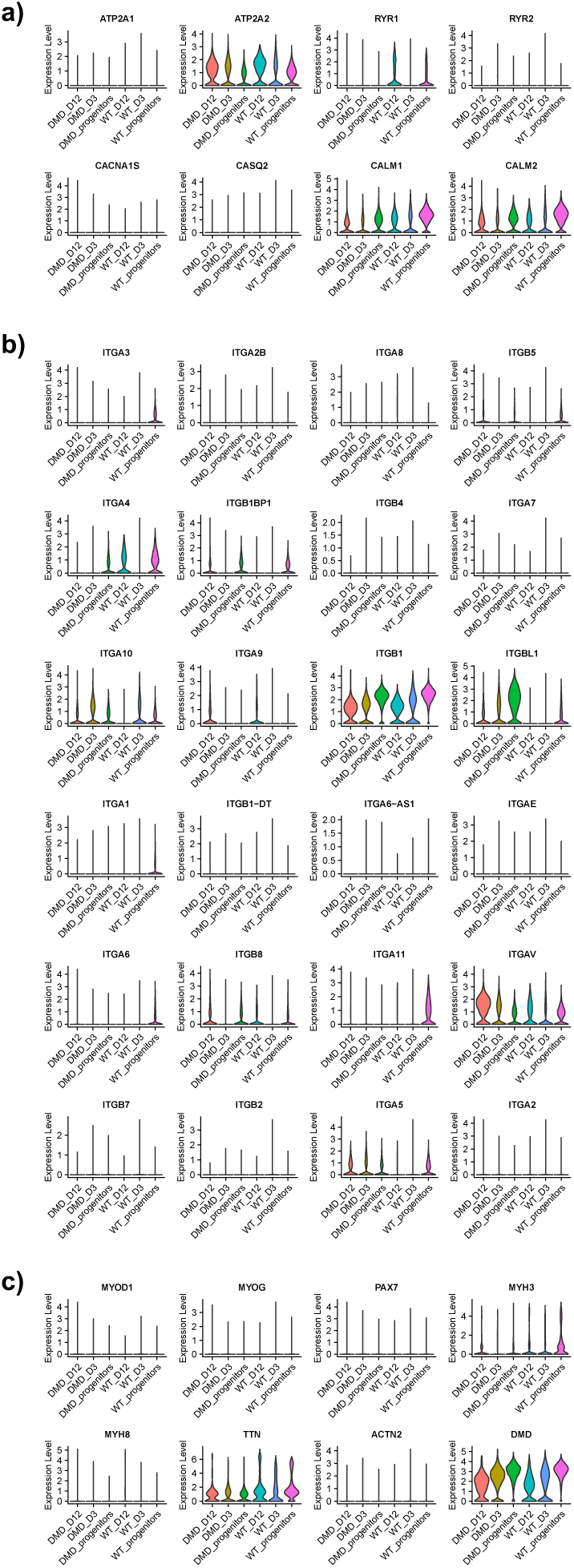
Expression levels of genes related to calcium handling (a), ITG genes (b) and myogenic makers (c) obtained from single nuclei RNA sequencing of UC3-4 (WT) and UC3-4 cl63 (DMD) myogenic progenitors as well as microtissue 3 and 12 days after casting.

